# Structural basis of UCUU RNA motif recognition by splicing factor RBM20

**DOI:** 10.1101/823351

**Authors:** Santosh Kumar Upadhyay, Cameron D. Mackereth

**Affiliations:** CSIR-Institute of Genomics and Integrative Biology, New Delhi 110020, India; Univ. Bordeaux, Institut Européen de Chimie et Biologie, 2 rue Robert Escarpit, 33607 Pessac Cedex, France; Inserm U1212, CNRS UMR5320, ARNA Laboratory, 146 rue Léo Saignat, 33076 Bordeaux Cedex, France

**Keywords:** cardiomyopathy, alternative splicing, NMR spectroscopy, RRM, RBM20, polypyrimidine tract-binding protein, RNA

## Abstract

The vertebrate splicing factor RBM20 (RNA Binding Motif protein 20) regulates protein isoforms important for heart development and function, with mutations in the gene linked to cardiomyopathy. Previous studies have identified the four base RNA motif UCUU as a common element in pre-mRNA targeted by RBM20. Here, we have determined the structure of the RNA Recognition Motif (RRM) domain from mouse RBM20 bound to RNA containing a UCUU sequence. The atomic details show that the RRM domain spans a larger region than initially proposed in order to interact with the complete UCUU motif, with a well-folded C-terminal helix encoded by exon 8 critical for high affinity binding. This helix only forms upon binding RNA with the final uracil, and removing the helix reduces affinity as well as specificity. We therefore find that RBM20 uses a coupled folding-binding mechanism by the C-terminal helix to specifically recognize the UCUU RNA motif.

## INTRODUCTION

Healthy cardiac development and function requires the regulated expression of many heart-specific genes. For several of these gene products, additional control through alternative splicing regulates a balance between cardiac protein isoforms that contain isoform-specific properties. A major example is the muscle protein titin, a giant protein that spans half the sarcomere and serves to modulate the elasticity of the stretched muscle (Maruyama et al., 1976; Wang et al., 1979; Azad et al., 2019). Titin exists as several isoforms and undergoes a notable shift in size from larger forms in early heart development towards shorter forms in the adult (reviews include Granzier and Labeit, 2002; Guo et al., 2010; Gigli et al., 2016). The functional outcome of these alternative isoforms is to change titin from the longer compliant form toward the shorter stiffer form, thus affecting passive muscle tension during the cardiac cycle. Disruption of this splicing regulation leads to abnormal ratios of the compliant and stiff forms of titin, resulting in heart disease. For example, failure to produce sufficient levels of the shorter titin isoform can lead to abnormally compliant titin in dilated cardiomyopathy (DCM).

In a search for factors that regulate titin splicing, a chromosomal deletion in *Rbm20* (RNA Binding Motif 20) was isolated as the genetic cause of titin mis-splicing from a mutant rat strain (Guo et al., 2012; Greaser et al., 2005). *RBM20* was itself first identified in a search for a familial genetic basis of DCM in human patients (Brauch et al., 2009). Subsequent investigation has identified additional patients and mutations involving *RBM20* related to DCM (Zhao et al., 2015b; Long et al., 2017; Wells et al., 2013; Waldmüller et al., 2015; Chami et al., 2014; Refaat et al., 2012; Rampersaud et al., 2011; Millat et al., 2011; Li et al., 2010; Robyns et al., 2019), as well as cardiac arrhythmia (Nielsen et al., 2018; Parikh et al., 2019), pediatric restrictive cardiomyopathy (Rindler et al., 2017) and left-ventricular non-compaction (Sedaghat-Hamedani et al., 2017). Although representing only 3 % of idiopathic cases, patients with DCM that have mutant RBM20 correlate with earlier disease onset, high penetrance, and requirement for heart transplantation (Brauch et al., 2009; Li et al., 2010; Kayvanpour et al., 2017; Wells et al., 2013; Hey et al., 2019).

Biological characterization of RBM20 has incorporated rat, mouse and *in vitro* studies. Expression of *Rbm20* is primarily localized to the heart, with lower expression in other striated muscle (Guo et al., 2012; Beraldi et al., 2014). Rats with homozygous or heterozygous loss of functional *Rbm20* mimic human DCM symptoms as well as defects in the heart structure, age-related fibrosis and less capacity for exercise (Guo et al., 2012; Guo et al., 2013). Loss of RBM20 function in rats also coincides with extensive deregulation of titin splice isoforms towards an abnormal form of titin with all exons retained (Greaser et al., 2005; Guo et al., 2012). Comparison to normal processing of titin pre-mRNA shows that RBM20 is required for several intron retention and exon skipping events, as well as the creation of circRNA (Li et al., 2013). Mice with homozygous deletion of *Rbm20* also reveal a large number of RBM20-dependent titin circRNA (Khan et al., 2016).

In addition to titin, RBM20 also regulates the splicing patterns of other transcripts that may contribute to the severity of DCM or promote separate cardiac pathologies. Human, rat and mouse studies identified additional splicing targets that encode proteins which bind and transport Ca^2+^ (Guo et al., 2012; van den Hoogenhof et al., 2018; Guo et al., 2018; Maatz et al., 2014; Wyles et al., 2016; Beraldi et al., 2014) such as the calcium channel ryanodine receptor 2 (*RYR2*), Ca^2+^/calmodulin-dependent protein kinase II delta (*CAMK2δ*), and the calcium channel voltage-gated L type alpha 1C subunit (*CACNA1C*/*Ca_v_1.2*). Other characterized targets include formin homology 2 domain containing 3 (*FHOD3*), Z-band alternatively spliced PDZ-motif protein (*ZASP*/*LDB3*/*CYPHER*), and the PDZ and LIM domain protein 5 (*PDLIM5*/*ENH*) (Maatz et al., 2014; Guo et al., 2012; Lorenzi et al., 2019; Ito et al., 2016).

The RBM20 protein sequence is largely devoid of identifiable structured regions except for the central RNA Recognition Motif (RRM) domain as well as two small putative zinc finger domains (ZnF) (Fig. 1A). To investigate a direct role in binding RNA, PAR-CLIP in HEK293 cells and HITS-CLIP in rat cardiomyocytes identified numerous transcripts that purified with tagged RBM20 (Maatz et al., 2014). Analysis of the sequences revealed a prominently conserved RNA motif composed of the tetramer UCUU: this motif is enriched 50 nucleotides before and 100 nucleotides after exons regulated by RBM20, which largely fall into the categories of mutually exclusive or cassette exons (Maatz et al., 2014). Furthermore, mutation of the UCUU motif to <u>CGCG</u> or <u>CG</u>UU sequences abolishes direct RBM20 binding to the *Ryr2* transcript, with reduction in binding by a single base change to UCU<u>G</u> (Maatz et al., 2014). Further analysis suggested that the RRM domain may indeed be responsible for UCUU recognition, since a construct from human RBM20 that covers the canonical RRM fold (residues 511-601) interacts only with titin-derived oligonucleotides that contain UCUU motifs (Dauksaite and Gotthardt, 2018). A minimal RRM region was also shown to bind to a titin transcript around titin exon 50 (Guo et al., 2012). Mouse strains with the RRM domain removed by deleting exons 6 and 7 of rbm20 result in mis-spliced titin, *Camk2d* and *Lbd3* (Methawasin et al., 2014) and increased titin stiffness in the diaphragm (van der Pijl et al., 2018). However, cardiac pathology in this strain was less pronounced than for a larger size *Rbm20* deletion (van den Hoogenhof et al., 2018). Although not corresponding to a folded domain, a region C-terminal to the RRM domain is enriched in arginine and serine residues (RS), and most of the disease mutations map to this small segment (Brauch et al., 2009) (Li et al., 2010; Watanabe et al., 2018). Improper phosphorylation of key serine residues in this RS domain prevents proper nuclear localization of RBM20 mutants (Murayama et al., 2018).

**Figure 1.**
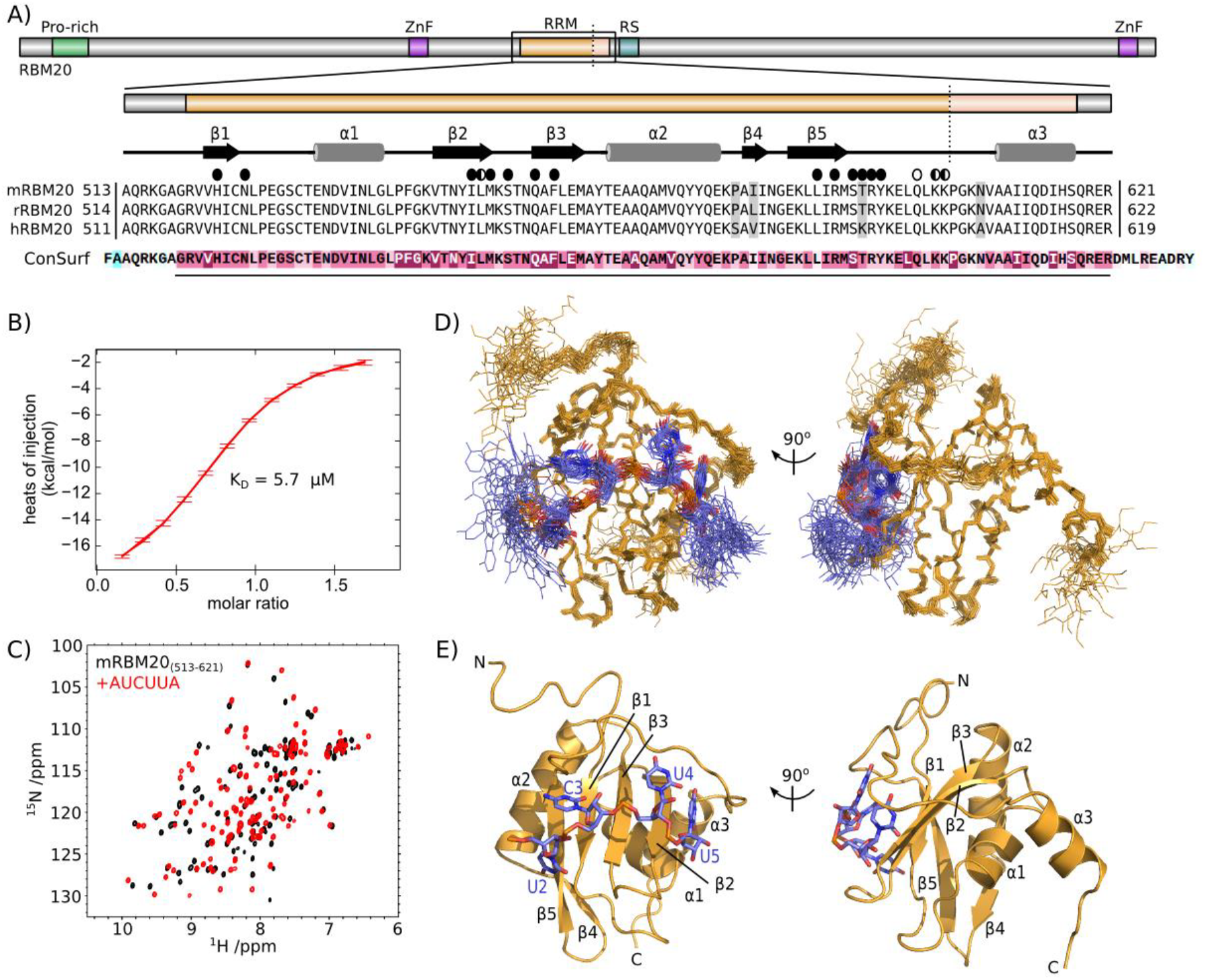
Recognition of UCUU by the RBM20 RRM domain. (**A**) Domain composition of RBM20 with expanded details for the RRM domain. The alignment is composed of RBM20 sequences from mouse (mRBM20, UniProt Q3UQS8), rat (rRBM20; UniProt E9PT37) and human (hRBM20; UniProt Q5T481). The vertical dotted line shows the C-terminal end of previous RRM domain constructs. Secondary structure elements from the NMR structure are indicated above the alignment. Residues that contact the RNA by their sidechain and backbone atoms are indicated by full and open circles, respectively. Results from the ConSurf analysis is shown below the alignment, with the region of high conservation indicated with a line, and coloured purple. (**B**) Representative isothermal titration calorimetry (ITC) data for mRBM20_(513-621)_ binding to AUCUUA RNA. (**C**) ^15^N-HSQC overlay of [^15^N]mRBM20_(513-621)_ in the absence (black) and presence (red) of 1.2 molar equivalents AUCUUA RNA. Annotated spectra can be found in **Supplementary Fig 1**. (**D**) Ensemble of 25 lowest energy structures calculated for mRBM20_(513-621)_ (backbone heavy atoms, orange lines) bound to AUCUUA RNA (all heavy atoms, purple lines). (**E**) Lowest energy structure model for the UCUU motif (all heavy atoms, purple sticks) and mRBM20 (residue 520-617, orange cartoon). RNA bases and protein secondary structure elements are labeled.

Given a biological role for the RRM domain and possible specificity for the UCUU motif exhibited by RBM20, we have used a structural approach to study RNA binding by the mouse RBM20 RRM domain. The atomic details confirm a role for the RRM domain in the direct interaction with the previously defined UCUU RNA motif. In addition to base recognition by the canonical RRM fold, we find that the specificity for the terminal uracil in the motif is coupled with folding of an extra C-terminal α-helix encoded by exon 8 of the *RBM20* gene.

## RESULTS

### RNA-binding by the RRM domain of RBM20

In order to first determine the boundaries of the construct to be produced for structural studies, we looked at sequence conservation around the RRM domain for RBM20 using ConSurf (Berezin et al., 2004; Ashkenazy et al., 2016) within the PredictProtein server (Yachdav et al., 2014) (Fig. 1A). The N-terminus of the RRM domain was clearly defined at the glycine preceding strand β1, whereas the C-terminal limit of conservation extended past the residues required for a typical RRM domain fold. To prevent unwanted truncation, we decided to include 18 additional residues at the C-terminus (Fig. 1A). This construct derived from mouse RBM20 (hereafter mRBM20_(513-621)_) produced soluble protein and displayed NMR spectroscopy data consistent with a folded domain (Fig. 1C, black spectrum; **Supplementary Figure 1**).

Our choice for RNA ligand was based on previous studies that have identified UCUU as an enriched RNA motif in RBM20-based PAR-CLIP of HEK293 cells and HITS-CLIP on rat cardiomyocytes (Maatz et al., 2014). In particular, the UCUU sequence in *Ryr2* is required for direct interaction by RBM20 (Maatz et al., 2014), and therefore we have selected a ligand based on this sequence. To help with synthesis and purification of the RNA oligonucleotide, we have included an extra adenine at the 5’ and 3’ ends of the motif derived from the *Ryr2* sequence. Isothermal titration calorimetry (ITC) using this AUCUUA RNA with mRBM20_(513-621)_ shows a 1:1 interaction with a K_D_ of 5.7 μM (Fig. 1B, Table 1). To provide additional details of binding specificity we performed a series of ITC measurements in which we made conservative mutations of the RNA ligand in which each base was mutated to the other purine or pyrimidine, respectively (Table 1). There were no significant changes in affinity upon mutation of the first or last base to guanine, in keeping with the primary importance of the central UCUU motif. Changing the first uracil of the motif to cytosine resulted in only a twofold increase in K_D_ (13 μM versus 5.7 μM). In contrast, mutation of each base in the remaining CUU sequence resulted in larger K_D_ increases of 16 to 36 times that of the original motif (K_D_ values of 95 to 210 μM).

**Table 1.**
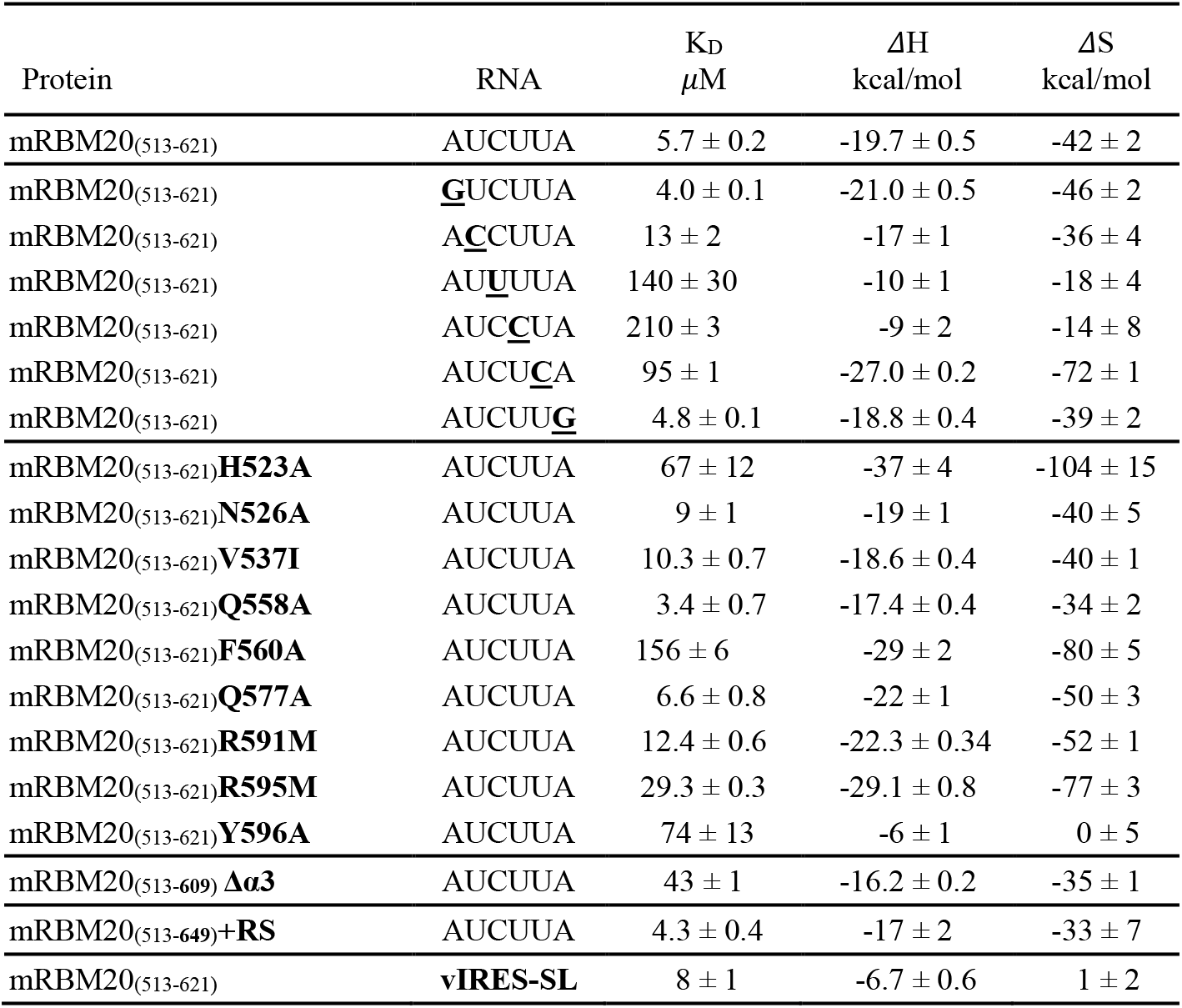
Isothermal titration calorimetry (ITC) measurements

Despite the relatively modest affinity to AUCUUA RNA, we found that addition of this ligand to ^15^N-labeled mRBM20_(513-621)_ resulted in clear backbone amide chemical shift perturbation for a majority of the residues (Fig. 1C, red spectrum). This effect is consistent with the formation of a stable and intimate protein-RNA complex, and in fact involves a greater number of residues than would be expected for the simple binding of the RNA to one face of the domain. Given the high quality of the resulting spectrum we decided to proceed directly to determine the atomic structure of the mRBM20 RRM domain bound to AUCUUA RNA.

### Molecular details of RNA binding to RBM20

Using a combination of distance, dihedral and residual dipolar coupling restraints, we determined an ensemble of 25 structures for mRBM20_(513-621)_ bound to AUCUUA RNA (Fig. 1D, Table 2). The most notable feature in the complex is the presence of an additional C-terminal helix (α3) that follows the canonical RRM fold (Fig. 1E). The atomic details also illustrate specific interaction between all four bases in the UCUU motif with either sidechain or backbone atoms of mRBM20_(513-621)_.

**Table 2.**
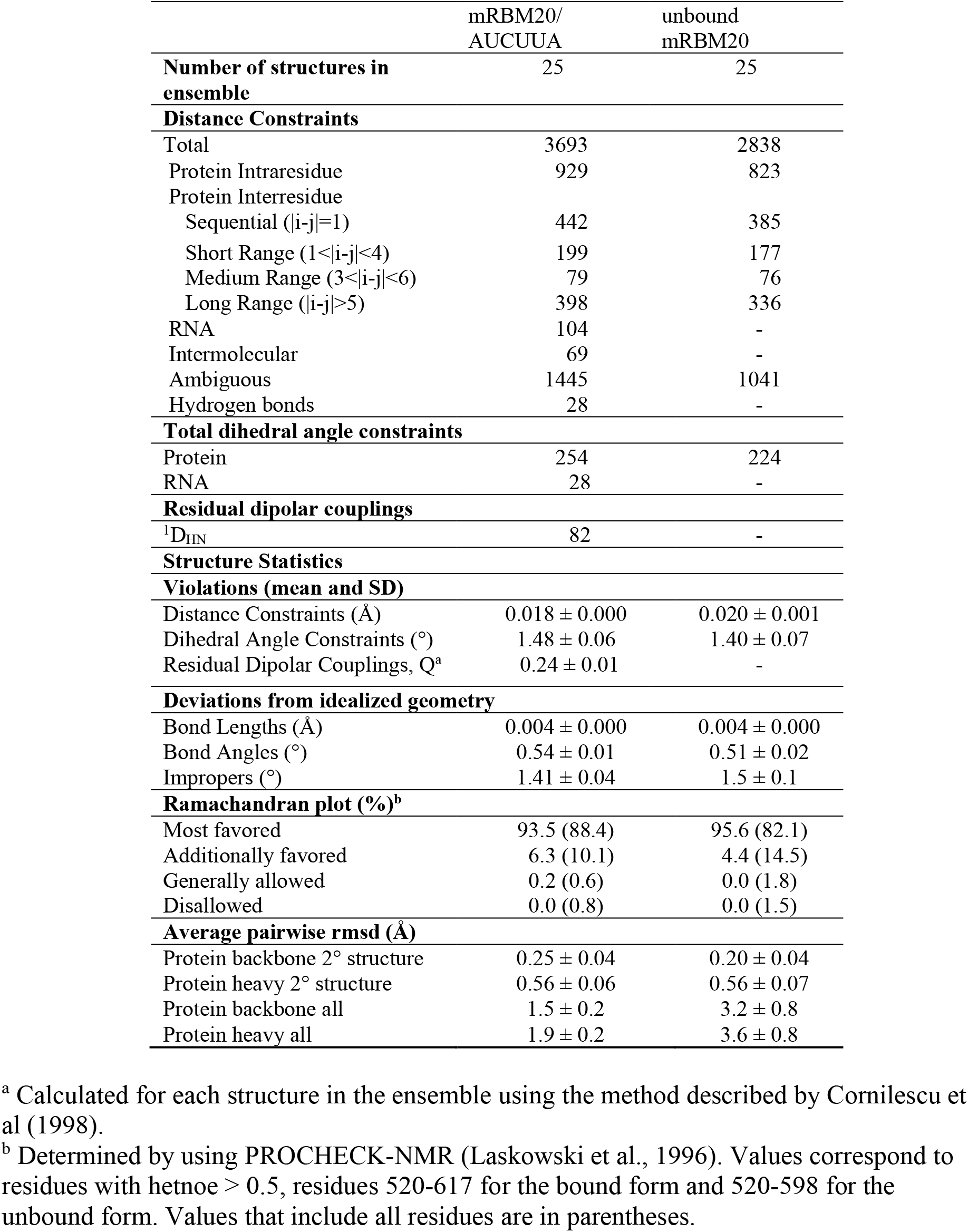
NMR and refinement statistics for the RNA-bound and unbound RBM20_(513-621)_

Starting with the first uracil, U2 in the structure, this base is located above Leu589 with hydrogen bonds to the side chain of Asn526, and further contact between Arg591 and the 5’ phosphate (Fig. 2A). Mutation of N526A results in only a small decrease in affinity (K_D_ of 9 μM) in keeping with the mild specificity for uracil. The sole cytosine, C3, stacks onto His523 and the mutation H523A reduces affinity by a factor of 12 (K_D_ of 67 μM; Table 1). The cytosine-specific amino group is recognized by a hydrogen bond from the sidechain of Thr594, with additional hydrogen bonds from Ser593 and the backbone amide of Thr594 (Fig. 2B). The 5’ phosphate is in contact with the sidechains of Gln558 and Arg595, but only mutation of Arg affects binding affinity (Table 1, K_D_ of 29.3 μM for R595M versus K_D_ of 3.4 μM for Q558A). The U4 base stacks onto Phe560, with mutation F560A resulting in an increased K_D_ of 156 μM (Table 1). In RBM20, there is notable binding affinity to the ribonucleotide granted by residues in the peptide linker that lies on top of the bound RNA. For U4, a pair of hydrogen bonds from the backbone atoms of Gln600 aids in uracil specificity (Fig. 2C). In addition, the aromatic ring of Tyr596, along with the sidechain of Phe560, defines a hydrophobic cleft which contacts hydrogens H5 and H6 of U4. Removal of the tyrosine aromatic ring in Y596A increases the K_D_ to 74 μM (Table 1).

**Figure 2.**
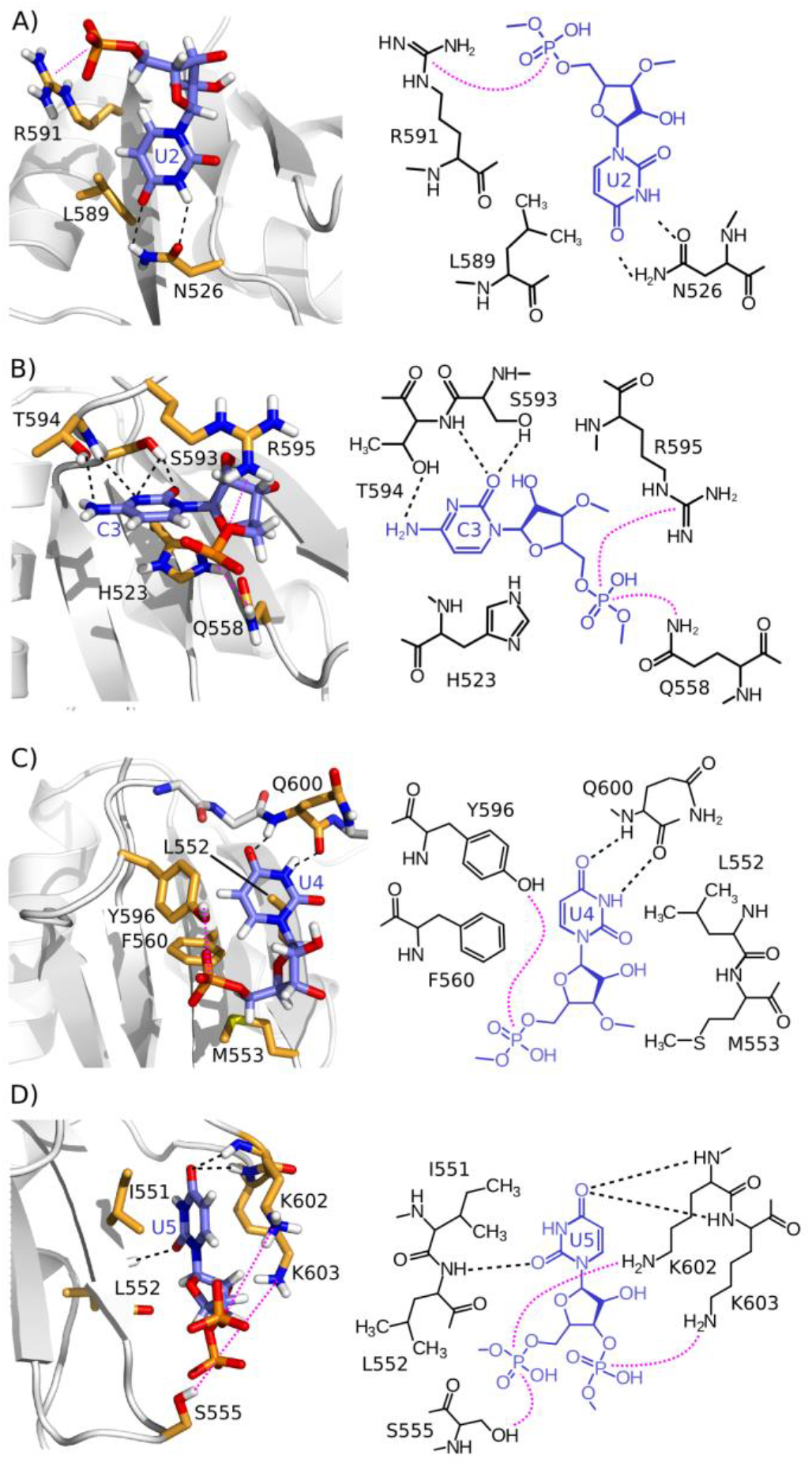
Molecular recognition of the UCUU motif by the RBM20 RRM domain. (**A-D**) Close up views of the intermolecular ^contacts between mRBM20_(513-621)_ and (**A**)^ U2, (**B**) C3, (**C**) U4, and (**D**) U5 of the UCUU motif.

Compared to the canonical recognition of C3 and U4 across the strands β1 and β3, the final uracil in the motif, U5, binds in an atypical position between strand β2 and the loop before the extra C-terminal helix (Fig. 2D). Hydrogen bonds from the backbone carbonyl of Leu552 and the backbone amides of Lys602 and Lys603 appear to guide specificity for uracil in this position. Due to the involvement of backbone atoms in this recognition, a simple site-directed mutation strategy can not be used to perturb binding of U5.

### The C-terminal helix is required for RNA binding

Based on the structure of the RNA-bound complex, the extra C-terminal helix α3 in mRBM20_(513-621)_ likely plays a role in the recognition of the 3’ uracil of the UCUU motif. By quantifying the change in ^1^H,^15^N amide crosspeak positions from the two spectra in Fig. 1C, it is evident that addition of AUCUUA RNA causes widespread perturbation throughout mRBM20_(513-621)_ (Fig. 3A). The changes caused by ligand binding naturally includes residues that lie below and above the bound RNA and are in direct contact with the ribonucleotides (Fig 3B). However, there is also significant chemical shift perturbation for all residues within the C-terminal helix, despite the fact that this helix lies distant to the bound RNA, on the opposite side of the RRM domain (Fig 3B).

**Figure 3.**
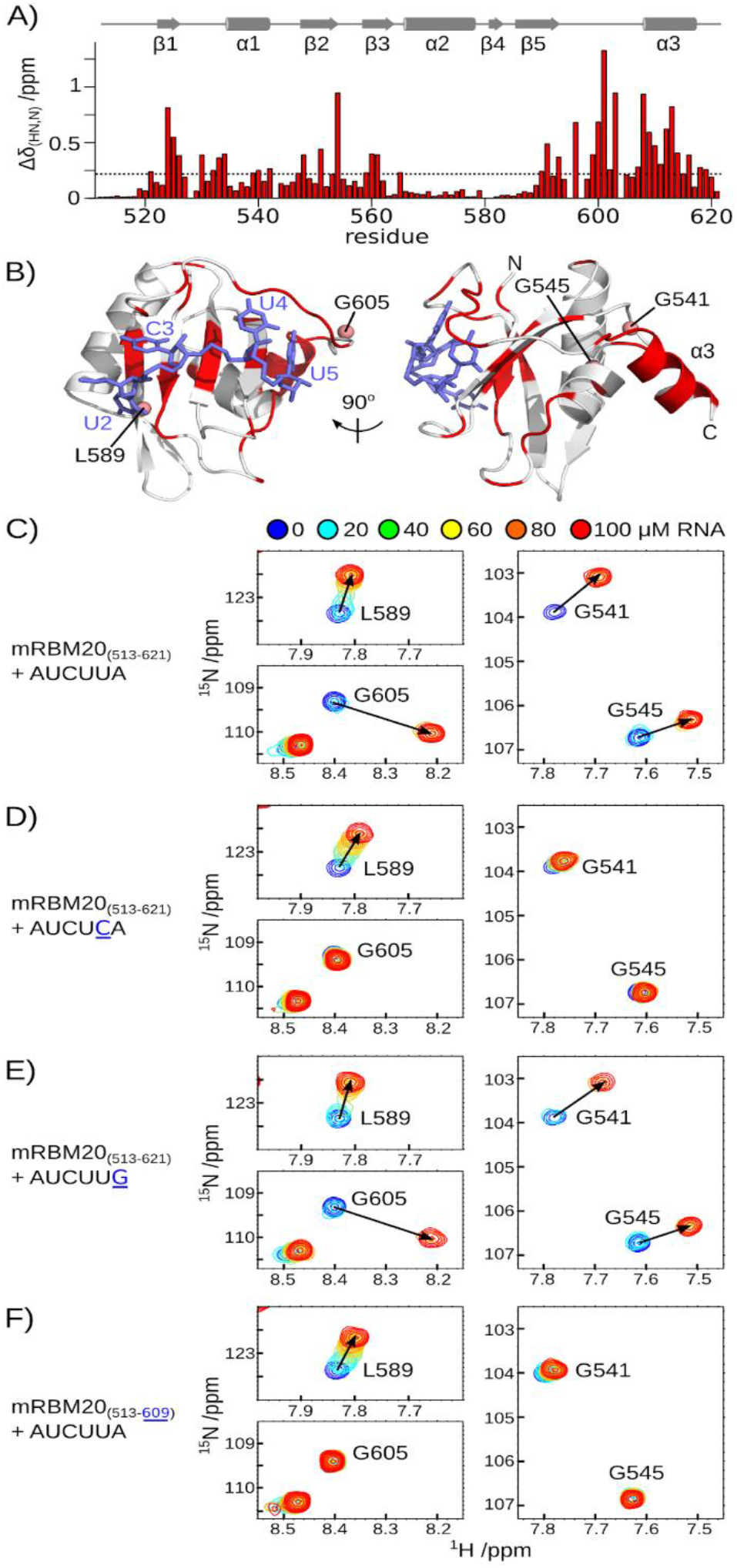
Helix α3 and nearby residues are affected by RNA binding. (**A**) Backbone amide chemical shift perturbation (Δδ_HN,N_) resulting from AUCUUA binding to mRBM20_(513-621)_ from the spectra shown in Fig. 1C, calculated as ((Δδ_HN_)^2+^(0.14*Δδ_N_)^2^)^0.5^. Secondary structure elements of RNA-bound mRBM20_(513-621)_ are shown above the histogram. (**B**) Residues with Δδ_HN,N_ greater than the average (0.23 ppm) are coloured red on the same views of RNA-bound mRBM20_(513-621)_ as shown in Fig. 1E. (**C-F**) Selected regions of the ^1^H,^15^N-HSQC spectra from 100 μM [^15^N]mRBM20_(513-621)_ titrated with 0, 20, 40, 60, 80 and 100 μM RNA. The locations of Leu589, Gly605, Gly541 and Gly545 backbone amides in mRBM20_(513-621)_ is indicated in (C). Complete spectra are shown in **Supplementary Figure 3**.

To gain further insight into the mechanism of RNA binding at the level of individual residues, we performed a series of titrations followed by NMR spectroscopy (**Supplementary Fig. 3**). Starting with mRBM20_(513-621)_, we followed the binding of the first and last uracil within AUCUUA RNA by changes in ^1^H,^15^N-HSQC crosspeak positions for Leu589 and Gly605, respectively (Fig. 3C). In addition, we find that residues distal to the RNA binding surface, but in contact with helix α3, are similarly affected by the ligand (Gly541 and Gly545; Fig. 3C).

From our initial characterization of RNA binding to mRBM20_(513-621)_ we noted that changing the terminal uracil to cytosine reduced binding affinity. When we followed titration of mRBM20_(513-621)_ with this AUCU<u>C</u>A RNA, the corresponding ^1^H,^15^N-HSQC show that RNA still induced changes in Leu589 (Fig. 3D). Therefore, the first uracil in AUCU<u>C</u>A is likely still recognized by RBM20. In contrast, the Gly605 crosspeak is unperturbed, implying that the introduced cytosine in position 5 is no longer able to interact with the protein (Fig. 3D). At the same time, the crosspeaks of Gly541 and Gly545 are also unaffected by RNA binding (Fig. 3D). Together, the data indicate that changing the terminal uracil to cytosine prevents interaction with mRBM20_(513-621)_ and that this loss in binding also prevents chemical shift perturbation for residues in the C-terminal helix. As a control, the titration of mRBM20_(513-621)_ with RNA ligand AUCUC<u>G</u>, in which the final adenine is replaced by guanine and affinity is maintained, does not affect the binding behaviour in the measured NMR spectra (Fig. 3E).

We next designed a truncated form, mRBM20_(513-609)_Δα3, in which only the α3 helical region has been removed, but all RNA-binding residues are retained. From ITC measurements, the loss of α3 results in a decreased affinity by a factor of eight (K_D_ of 43 μM; Table 1), supporting a key role for this helix in RNA recognition. When the mRBM20_(513-609)_Δα3 mutant is titrated with AUCUUA RNA, the corresponding ^1^H,^15^N-HSQC NMR data show that the overall domain fold is retained, and that Leu589 still reports on binding by the first uracil (Fig. 3F). In contrast, the Gly605 crosspeak is unperturbed, implying that there is no interaction between the protein and the final base in the RNA motif even though the preferred uracil is present (Fig. 3F). This lack of perturbation once again extends to the crosspeaks of Gly541 and Gly545 (Fig. 3F). The results indicate that although the C-terminal helix does not directly interact with the RNA, the helix is nonetheless required to form the binding site for the final uracil in the motif. In addition, the RNA-induced chemical shift changes observed for Gly541 and Gly545 requires the presence of the C-terminal helix. We therefore hypothesize that helix α3 may not be a constant part of the RBM20 RRM domain, but may be selectively stabilized only when the final uracil is present in the RNA motif.

### Disordered C-terminus in the unbound state

To determine the nature of helix α3 in the unbound state, we calculated an ensemble of NMR-based structures for the free mRBM20_(513-621)_ (Fig. 4A, Table 2). The core RRM fold is nearly identical between the bound and free states with backbone rmsd of 0.01 Å for residues 520-598 calculated between the two ensembles using SuperPose (Maiti et al., 2004). In contrast, the C-terminal region in unbound mRBM20_(513-621)_ clearly lacks a stable helical fold (Fig. 4B). This region instead displays increased disorder as compared to the RNA-bound state, and by NMR spectroscopy is more flexible (Fig. 4C). A small plateau of similar backbone dynamics for a stretch of residues in the C-terminus might correspond to a transient helix for the unbound mRBM20_(513-621)_. A very slight helicity in the α3 region in the free state is also indicated upon analysis of backbone ^13^Cα secondary chemical shifts (Fig. 4D). Upon binding RNA, the helix α3 region and preceding loop shifts towards less dynamics to form the RNA-bound complex.

**Figure 4.**
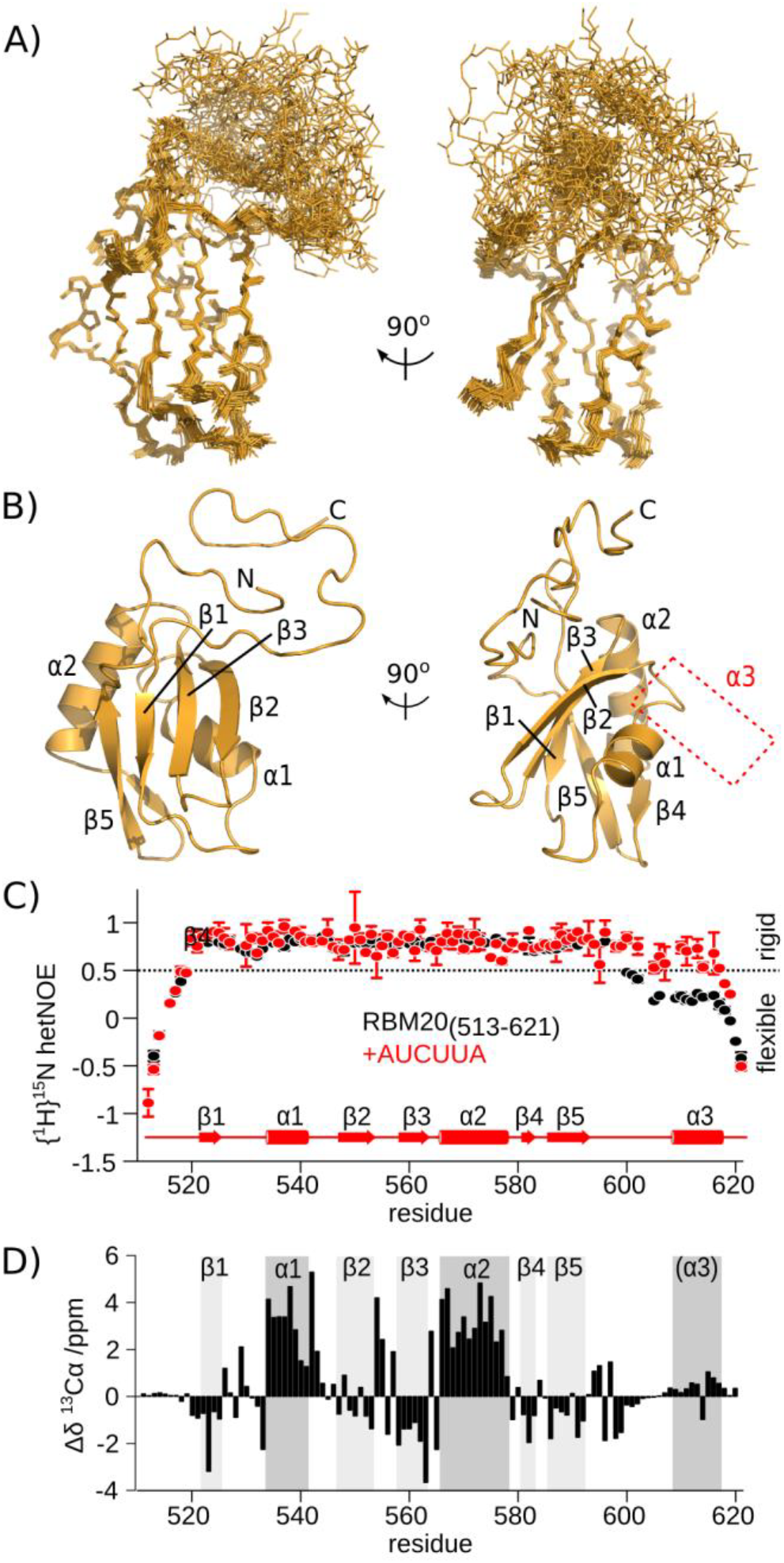
The helix α3 region is disordered in unbound mRBM20_(513-621)_. (**A**) Ensemble of 25 lowest energy structures calculated for unbound mRBM20_(513-621)_ (backbone heavy atoms, orange lines). (**B**) Lowest energy structure model for mRBM20 (orange cartoon). Protein secondary structure elements are labeled. The location of α3 in the RNA-bound complex is also indicated. (**C**) Flexibility of the backbone amides as measured by {^1^H}^15^N hetNOE of the unbound (black) and RNA-bound (red) [^15^N]mRBM20_(513-621)_. The secondary structure elements of RNA-bound mRBM20_(513-621)_ is also shown. Residues with {^1^H}^15^N hetNOE values less than 0.5 are considered to be disordered. (D) ^13^Cα secondary chemical shift (Δδ) of the unbound mRBM20_(513-621)_ compared to a disordered peptide as predicted by using ncIDP (Tamiola et al., 2010). The secondary structure elements of RNA-bound mRBM20_(513-621)_ are indicated by grey boxes.

### Residues implicated in disease have little effect on RNA binding

Most of the *RBM20* mutations implicated in cardiomyopathy are localized within the RS region, and these mutants mainly disrupt nuclear localization (Murayama et al., 2018). To see if the addition of the RS region could in general affect RNA binding, we prepared a construct with the C-terminus extended to residue 649. This longer construct had the same affinity to AUCUUA RNA as mRBM20_(531-621)_ (K_D_ of 4.3 μM compared to 5.7 μM; Table 1). The similar K_D_ values rule out a significant contribution to RNA binding by the RS region, at least for the unphosphorylated protein, and therefore mutations in this area would likely not act via simple changes in RNA binding. The only RRM domain mutant with possible link to disease, V537I, is located distant to the RNA-binding surface but is close to the area contacted by the C-terminal helix. By ITC, the affinity of V537I to AUCUUA RNA shows a modest two-fold reduction (K_D_ of 10.3 μM; Table 1).

### Structural similarity to polypyrimidine tract-binding protein

Given the importance of the added C-terminal helix in the RBM20 RRM domain, we searched other RRM domains that may use the same RNA-binding mechanism. The only candidate we found was the first RRM domain of polypyrimidine tract-binding protein (PTBP1/PTB4) bound to a viral internal ribosome entry site (vIRES) stemloop RNA (PDB ID 2N3O; (Dorn et al., 2017). A sequence similarity was also previously identified in the RNP1 motifs of RBM20 and RRM1 of PTBP1 (Filippello et al., 2013). In both RNA-bound structures, a UCUU sequence is involved in the interaction with a highly conserved hydrogen bond network to the CUU bases (Fig. 5A). In particular, the 3’ uracil in both cases is recognized by hydrogen bonds to backbone amides in the loop preceding helix α3 (Fig. 5A). Despite the presence of α3 in both structures, the sequence conservation is low in the region C-terminal to the core RRM fold (Fig. 5B,**C**). Most residues that lie atop the bound RNA differ between RBM20 and PTBP1 RRM1 (Fig. 5C, left). More noticeable is that the C-terminal helix is significantly shifted between the two structures, although the overall path of the C-terminal residues is similar (Fig. 5C, right). In terms of the RNA ligand, a difference between the two complexes is that RBM20 is bound to a short single-stranded RNA, whereas PTBP1 RRM1 binds UCUU within the loop of the vIRES stemloop RNA. In the vIRES stemloop the first uracil is base-paired within the stem structure, and thus cannot be bound in the same orientation as in the ssRNA ligand. Using ITC we nevertheless find that mRBM20_(513-621)_ can also interact with the vIRES stemloop UCUU with almost the same affinity as for ssRNA (K_D_ of 8 μM verses 5.7 μM; Table 1).

**Figure 5.**
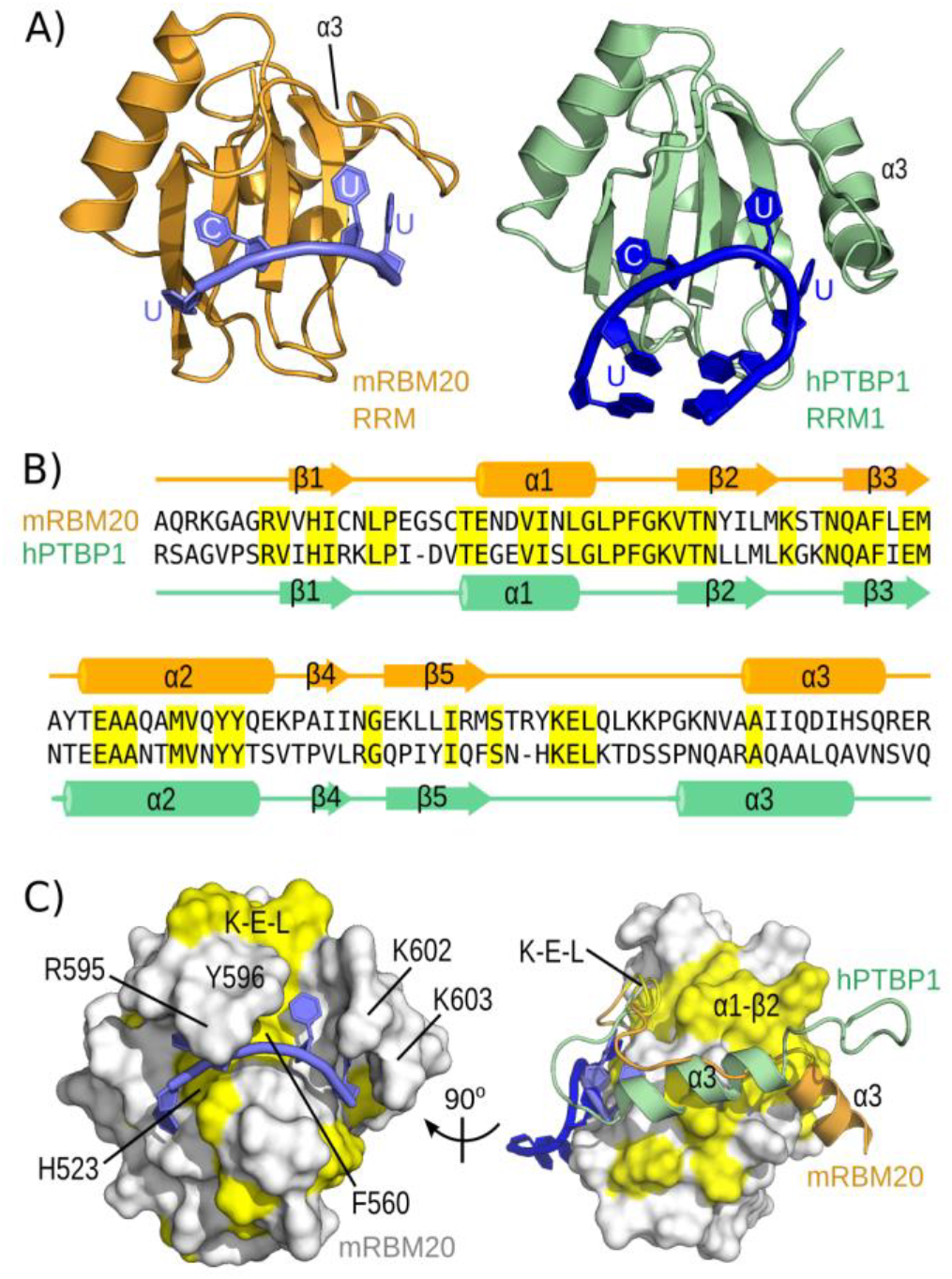
Structural similarity to RNA-bound PTBP1 RRM1. (**A**) Cartoon representation of protein-RNA complexes involving mRBM20 RRM domain (left) or hPTBP1 RRM1 (right, from PDB ID 2N30) bound to RNA containing a UCUU motif. (**B**) Sequence alignment of mouse RBM20 RRM and human PTBP1 RRM1, with identical residues highlighted in yellow. Protein secondary structure elements are shown from each RNA-bound complex. (**C**) Conserved residues from (B) are coloured yellow on the model of RNA-bound mRBM20_(513-621)_ using the same views as in Fig. 1E. On the right, the C-terminal residues of both mRBM20 (orange) and hPTBP1 (light green) are shown in cartoon representation, with the core RRM domain fold as a surface representation.

## DISCUSSION

We have demonstrated in atomic detail the way in which the RRM domain from RBM20 is specific for the UCUU RNA sequence. Each base in the motif is recognized by the RRM domain, with an atypical binding of the final uracil coupled to stabilization of a C-terminal helix (Fig. 6A). These results establish the RRM domain as a major determinant of the RNA binding specificity of RBM20, and a basis of UCUU enrichment in target pre-mRNA.

**Figure 6.**
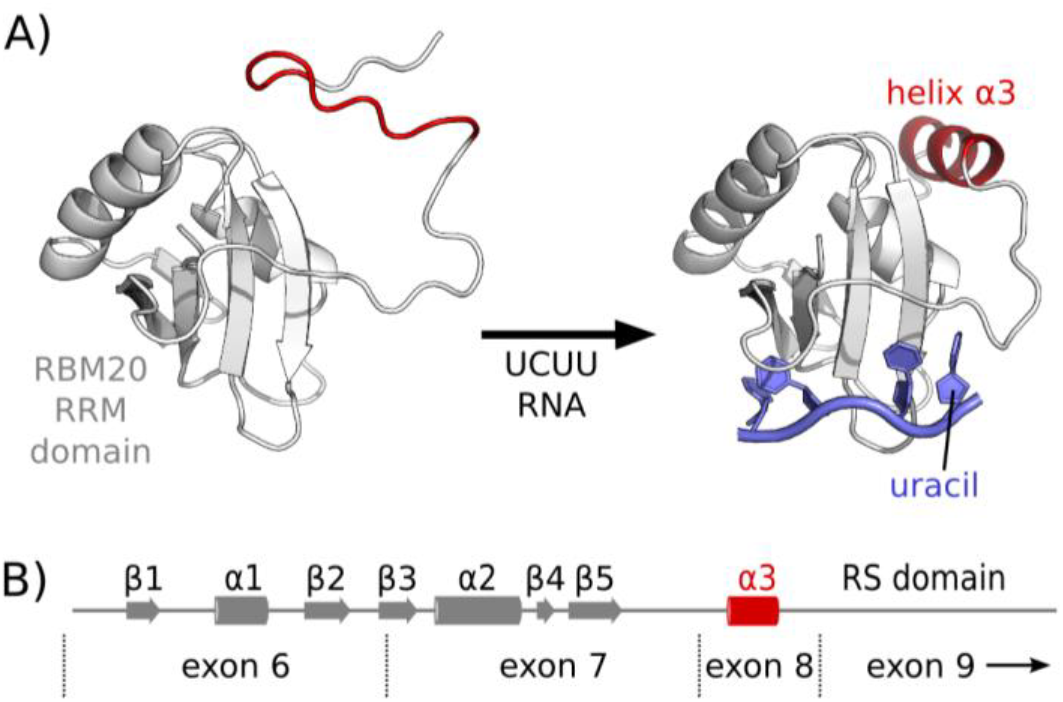
Model of RNA binding by RBM20. (A) In the unbound state, the RRM domain from RBM20 has a disordered C-terminus. Upon binding with an RNA ligand containing the sequence UCUU, the 3’ uracil combines with formation of a C-terminal helix to stabilize the protein-RNA complex. (C) The C-terminal α3 helix is encoded by exon 8 of the *RBM20* gene, in between exons 6 and 7 that encode the canonical RRM domain fold, and the RS domain in exon 9.

The main finding from the structural studies is the unusual role of the C-terminal helix in RNA recognition. Despite its importance in RNA binding, helix α3 itself does not directly interact with the bound uracil, but instead packs against β2 and helix α1 of the core RRM domain. This finding is notable since the low affinity binding by truncated construct mRBM20_(513-609)_Δα3, in which this helix is removed, nonetheless retains all of the residues and backbone atoms that directly contact RNA. In this context, the helix mainly represents an additional but key point of stabilization for the loop residues that directly contact RNA. This model explains the lack of sequence conservation between the C-terminal helices of RBM20 and PTBP1 RRM1 (Fig. 5B).

An absolute need for residues C-terminal to the canonical RRM fold means that exon 8 of RBM20 is a required part of the functional RRM domain. The complete RNA-binding region from *RBM20* therefore consists of the segment stretching from exon 6 to exon 8 (Fig. 6B), followed directly by the RS domain encoded by exon 9. This situation is similar to PTBP1 RRM1, in which the C-terminal helix is also encoded by the exon that follows the canonical RRM domain.

The C-terminal helix is critical to RNA selectivity, but the reverse is also true: binding of a high affinity UCUU motif is required to form the C-terminal helix. As a consequence, only by binding the complete motif would the residues following the canonical RRM domain condense into a stabilized helix. This triggered effect would induce lowered flexibility of the C-terminal residues upon binding the correct RNA sequence, and thus reduce the distance to the following domain such as the RS region in RBM20 or RRM2 in PTBP1. The RNA-triggered helix formation would also drastically alter the accessible surface features, and thus could either create or inhibit interaction with auxiliary binding partners.

It is anticipated that additional domains in RBM20, as well as interaction with other proteins, help to further restrict RNA binding to specific motifs in target pre-mRNA. For example, the two zinc finger domains in RBM20 may contribute directly to RNA binding. The C-terminal zinc finger is essential for RBM20 function (Dauksaite and Gotthardt, 2018), however so far neither of the two zinc fingers have been shown to bind RNA. The RS domain is required for nuclear localization (Murayama et al., 2018) and likely mediates protein-protein interactions (Maatz et al., 2014). We have shown that inclusion of the unphosphorylated RS domain does not directly affect affinity to AUCUUA RNA (Table 1).

In terms of protein-protein interactions, polypyrimidine tract-binding protein PTBP1 has been found to co-regulate splicing with RBM20 for *FHOD3* pre-mRNA (Lorenzi et al., 2019), and also for a mini-gene reporter based on titin (Dauksaite and Gotthardt, 2018). Given the similarity in UCUU binding mechanisms between RBM20 and RRM1 of PTBP1 (Fig. 5), this co-regulation could also involve some level of competition for shared binding sites. From our studies, RBM20 can equally recognize the UCUU motif in short single-stranded RNA as well as in the context of a loop in stemloop RNA (Table 1). However, the overall affinity for both is relatively moderate (K_D_ values of 5.7 and 8 μM, respectively). It is possible that RRM1 from PTBP1 exhibits a preference for UCUU in a particular context (Dorn et al., 2017), in which case PTBP1 binding would relegate RBM20 binding to the remaining UCUU sites. Polypyrimidine tract-binding protein contains three additional RRM domains that also favour UC-rich sequences (Oberstrass et al., 2005), and this intramolecular cooperativity may also function in competition with RBM20.

Another protein partner of RBM20 is RBM24, and these two factors co-regulate inclusion of exon 11 from *Enh* pre-mRNA (Ito et al., 2016). A direct interaction was found *in vitro* between the full-length RBM20 and RBM24 proteins, and a cooperative binding by both proteins might help define target specificity to the region upstream of exon 11. In support of functional cooperativity, single mouse knockouts of RBM20 or RBM24 have only minimal effect on exon 11, whereas simultaneously increasing or decreasing RBM20 and RBM24 shifts *Enh* towards the long or short isoform, respectively (Ito et al., 2016).

Additional insight into the function of RBM20 derives from the study of titin pre-mRNA processing. Extensive characterization of RBM20-dependent titin isoforms describes exon skipping, intron retention and the creation of circRNA (Li et al., 2013). Visualization of RBM20 in cardiomyocytes and muscle tissue show an organization into two nuclear clusters that colocalize with titin mRNA but not with other nuclear bodies, including paraspeckles or nuclear speckles (Li et al., 2013). Purification of RBM20-bound titin RNA shows that introns remain, and thus it has been proposed that these clusters represent intermediate or co-transcriptional steps in titin pre-mRNA processing during which RBM20 remains bound to extended regions of titin pre-mRNA (Li et al., 2013). Recent findings suggest that these RBM20 processing clusters may also connect chromosomal regions for other splice targets such as *CACNA1C* and *CAMK2D* (Bertero et al., 2019). Our molecular study of RNA binding by the RBM20 RRM domain helps in the understanding of binding site specificity, but the long residence time of RBM20 on pre-mRNA targets such as titin suggests the contribution of additional elements that stabilize the protein-RNA complexes. These additional contacts could include regions of RBM20 that mediate RBM20-RBM20 contacts, as well as possible contacts to proteins such as PTBP1 and RBM24.

Removal of the RRM domain in mouse RBM20 leads to a phenotype consistent with, but less severe than, strains in which the complete RBM20 protein function is abolished (Methawasin et al., 2014; Guo et al., 2012). Nevertheless, few *RBM20* disease mutations have so far been found within the RRM domain: there is a single example of a sporadic V535I mutation (V537I in mice) that has only limited effect on RNA binding (Table 1), and the proximal I536T mutation that may be secondary to mRNA processing defects in *LDB3* (Yamamoto et al., 2019). In contrast, the RS domain remains the hotspot of *RBM20* disease mutations, and mutations such as S637A can act as a dominant negative toward titin splicing (Murayama et al., 2018; Kimura, 2016). It may be that functional mutants of RBM20 that can still bind to native pre-mRNA binding sites are more deleterious that RBM20 mutant proteins that simply affect RNA-binding specificity. On the other hand, the modest effects observed for *Rbm20ΔRRM* mice illustrate putative benefits of adjusting RBM20 function to help mediate certain heart pathologies. Mice strains that model abnormal titin stiffness show that crosses with *Rbm20ΔRRM* mice help promote a more compliant titin in the offspring with improvement of muscle function (Methawasin et al., 2014; Buck et al., 2014). Similarly, abnormal diastolic function in mouse models of heart failure with preserved ejection fraction (HFpEF) can be largely restored with heterozygous expression of *Rbm20ΔRRM* (Hinze et al., 2016; Bull et al., 2016; Methawasin et al., 2016). In both cases, it is likely that effects on additional targets related to cardiac performance may contribute to phenotypic changes in the *Rbm20ΔRRM* mice, notably splicing and expression levels of proteins involved in calcium sensitivity and contraction regulation (Pulcastro et al., 2016; Guo et al., 2018).

The results obtained through mouse genetics provide support for therapeutic reduction of RBM20 function to help related human pathologies. Indirect modulation of RBM20, such as through thyroid hormone signalling (Zhu et al., 2015), the mTor pathway (Zhu et al., 2017) or by angiotensin II (Cai et al., 2019) could affect RBM20 function. Recently, several cardenolide compounds were discovered that target the RBM20-dependent regulation of titin splicing (Liss et al., 2018), however the mechanism did not appear to alter RNA-binding by the RRM domain. A direct targeting of RNA-binding based on the atomic details of the protein-RNA complex could serve as an additional therapeutic strategy.

## MATERIALS AND METHODS

### Cloning

Constructs containing the RRM domain of murine RBM20 were created from a cDNA provided by Pamela Lorenzi (University of Verona, Italy), together with PCR primers containing an *Nco*I restriction site in the forward primer, with a stop codon and *Acc65*I restriction site in the reverse primers. The primer details are included in **Supplementary Table 1**. The PCR products, as well as a modified pET-9d vector, were digested with *Nco*I and *Acc65*I, followed by ligation, to produce plasmids that encode constructs with an N-terminal hexahistidine (His_6_) tag and cleavage site for tobacco etch virus (TEV). The ligation products were transformed into *E. coli* DH5α and resulting plasmids were verified by sequencing. Mutants were created by performing an initial PCR step with internal forward and reverse primers that harbour the mutant sequence.

### Protein expression

*E. coli* BL21(DE3) *pLysY* (New England Biolabs) were transformed with plasmids encoding the various mRBM20 constructs. Subsequent colonies were used for initial overnight culture growth at 37 °C in lysogeny broth (LB) supplemented with 50 μg/ml kanamycin. Bacteria from the overnight cultures were used to start 500 mL cultures in LB for natural abundance protein, or in M9 minimal medium supplemented with isotopically-enriched compounds. For ^15^N-labeled protein, 1 g/L ^15^NH_4_Cl was added to the media, and additionally 2 g/L ^13^C-glucose was added for ^13^C,^15^N-labeled protein. For stereospecific assignment of methyl groups, the media was only enriched to 10% (w/w) ^13^C-glucose. Deuterated protein was grown in 99 % (v/v) ^2^H_2_O with 2 g/L ^2^H,^13^C-glucose and 1 g/l ^15^NH_4_Cl. In all cases, initial growth of the 500 mL cultures at 37 °C was followed by induction at an OD_600nm_ of 0.6 with 0.25 mM isopropyl β-D-1-thiogalactopyranoside (IPTG), and protein expression continued for 16 h at 25 °C. Cells were harvested by centrifugation at 4500 x *g* for 20 min at 4°C, resuspended in lysis buffer containing 5 mM imidazole, 50 mM Tris (pH 7.5), 500 mM NaCl, 5 % (v/v) glycerol, and stored at −80 °C in the presence of added lysozyme.

### Protein purification

Cells frozen in lysis buffer containing lysozyme were thawed on ice, with lysis aided by sonication. Soluble protein was separated from cellular debris by centrifugation at 20000 x *g* for 30 min at 4 °C. The supernatant was filtered through a GD/X 0.7 μm filter (GE Healthcare Life Sciences) and loaded onto 2 mL Nuvia IMAC Ni-charged resin (Bio-Rad Laboratories). The resin was then washed with 10 column volumes of buffer containing 5 mM imidazole, 50 mM Tris (pH 7.5), 500 mM NaCl, 5 % (v/v) glycerol followed by 5 volumes of the same buffer but with 25 mM imidazole. Protein elution used the same buffer with 500 mM imidazole. Fractions containing protein were pooled and exchanged to the initial buffer containing 5 mM imidazole by using a PD10 desalting column (GE Healthcare Life Sciences). His-tagged TEV protease (0.1 mg/ml final concentration) was added for overnight cleavage at 4 °C. The protease, hexahistidine tag and any uncleaved protein was removed by a second passage through the Nuvia IMAC Ni-charged resin. The purified samples were concentrated with 3 kDa MWCO Vivaspin centrifugal concentrators (Merck Millipore Corporation) followed by overnight dialysis in 20 mM sodium phosphate (pH 6.5) and 50 mM NaCl. Samples for NMR spectroscopy were supplemented with 2 mM dithiothreitol (DTT) and 10 % (v/v) ^2^H_2_O. Samples for Isothermal Titration Calorimetry (ITC) included 2.5 mM Tris(2-carboxyethyl)phosphine (TCEP) in the dialysis buffer. Protein purity was checked by SDS-PAGE and protein concentration was determined by absorbance at 280 nm with extinction coefficients obtained using ProtParam (http://web.expasy.org/protparam).

### RNA synthesis

RNA oligonucleotides were synthesized on an Expedite 8909 (PerSeptive Biosystems) from phosphoramidite monomers, and purified from a mix of butanol and water. AUCUUA RNA was also purchased (Sigma-Aldrich). RNA concentrations were determined by absorbance at 260 nm with extinction coefficients obtained from OligoAnalzer (https://eu.idtdna.com/calc/analyzer).

### NMR spectroscopy

NMR spectra were recorded at 298 K using a Bruker Neo spectrometer at 700 MHz or 800 MHz, equipped with a standard triple resonance gradient probe or cryoprobe, respectively. Bruker TopSpin versions 4.0 (Bruker BioSpin) was used to collect data. NMR data were processed with NMR Pipe/Draw (Delaglio et al., 1995) and analysed with Sparky 3 (T.D. Goddard and D.G. Kneller, University of California).

### Chemical shift assignment

Backbone ^1^H^N^, ^1^H^α^, ^13^C^α^, ^13^C^β^, ^13^C’ and ^15^N^H^ chemical shifts for mRBM20_(513-621)_ bound to AUCUUA were assigned based on 2D ^1^H,^15^N-HSQC, 3D ^15^N-HNCO, 3D ^15^N-HNCACO, 3D ^15^N-HNCA, 3D ^15^N-HNCACB, 3D ^15^N-CBCACONH, and 3D ^15^N-HNHA spectra. Aliphatic side chain protons were assigned based on 2D ^1^H,^13^C-HSQC, 3D C(CO)NH-TOCSY, 3D H(C)CH-TOCSY and 3D (H)CCH-TOCSY. Stereospecific assignment of leucine and valine methyl groups used a 10 % (w/w) ^13^C-glucose sample and a constant time ^1^H,^13^C-HSQC (Senn et al., 1989). Assignment of sidechain asparagine δ2 and glutamine ε2 amides used a 3D ^15^N-HSQC-NOESY (120 ms mixing time). Aromatic ^1^H and ^13^C nuclei were assigned based on 2D ^1^H,^13^C-HSQC and 3D ^13^C-HSQC-NOESY (150 ms mixing time). Non-exchanging RNA ^1^H nuclei were assigned in 99 % (v/v) D_2_O from natural abundance AUCUUA RNA in complex with [^2^H-99%]mRBM20_(513-621)_ by using 2D ^1^H,^1^H-NOESY (120 ms mixing time) and ^1^H,^1^H-TOCSY spectra. Exchanging RNA ^1^H nuclei were assigned by using a x2-filtered 2D ^1^H,^1^H-NOESY (150 ms mixing time) at 278 K on a sample of [^13^C,^15^N]mRBM20_(513-621)_ bound to AUCUUA.

For the unbound mRBM20_(513-621)_ the backbone ^1^H^N^, ^1^H^α^, ^13^C^α^, ^13^C^β^, ^13^C’ and ^15^N^H^ chemical shifts were assigned based on 2D ^1^H,^15^N-HSQC, 3D ^15^N-HNCO, 3D ^15^N-HNCACO, 3D ^15^N-HNCA, 3D ^15^N-HNCACB, and 3D ^15^N-CBCACONH spectra. Aliphatic side chain protons were assigned based on 2D ^1^H,^13^C-HSQC, 3D H(C)(CO)NH-TOCSY, 3D (H)C(CO)NH-TOCSY, 3D H(C)CH-TOCSY and 3D (H)CCH-TOCSY. Stereospecific assignment of leucine and valine methyl groups used a 10 % (w/w) ^13^C-glucose sample and a constant time ^1^H,^13^C-HSQC (Senn et al., 1989). Assignment of sidechain asparagine δ2 amdies and glutamine ε2 amides used a 3D ^15^N-HSQC-NOESY (120 ms mixing time). Aromatic ^1^H and ^13^C nuclei were assigned based on 2D ^1^H,^13^C-HSQC (^13^C offset 120 ppm) and 3D ^13^C-HSQC-NOESY (120 ms mixing time, ^13^C offset 125 ppm).

### Structure calculation

Structure ensembles were calculated by using Aria 2.3/CNS1.2 (Rieping et al., 2007) (Brunger, 2007), with final ensembles refined in explicit water and consisting of the 25 lowest energy structures from total of 100 calculated models. Complete refinement statistics are presented in Table 1.

For the RNA-bound complex, the majority of protein ^1^H distances were obtained using NOE crosspeaks from 3D ^15^N-HSQC-NOESY (120 ms mixing time), 3D ^13^C-HSQC-NOESY (150 ms mixing time), and 2D ^1^H,^1^H-NOESY (120 ms mixing time) spectra. RNA-RNA ^1^H distances were derived from a 2D ^1^H,^1^H-NOESY (120 ms mixing time) spectrum using natural abundance AUCUUA RNA in complex with a fully deuterated mRBM20_(513-621)_ in 100 % D_2_O.

Intramolecular distances were mostly derived from a x2-filtered 2D ^1^H,^1^H-NOESY spectrum (240 ms mixing time) on a sample of [^13^C,^15^N]mRBM20_(513-621)_ bound to AUCUUA, and a similar spectrum acquired at 278 K was used to identify NOE crosspeaks involving exchangeable ^1^H RNA nuclei. Protein dihedral angles were obtained by using TALOS-N (Shen and Bax, 2015) and SideR (Hansen et al., 2010a; Hansen et al., 2010b). RNA dihedral angles were based on TOCSY and NOESY crosspeak patterns. Following a preliminary structure calculation, residues within the centre of each α-helix were further restrained by hydrogen-bond restraints. Starting at iteration four, residual dipolar coupling (RDC) values were included based on alignment using the stretched-gel approach (Liu and Prestegard, 2010) by measuring interleaved spin state-selective TROSY experiments on RNA-bound [^15^N]-mRBM20_(513-621)_. For the aligned sample, a solution of the complex was added to a 1 cm cylinder of dried 5% 19:1 acrylamide:bisacrylamide and stretched into an NMR tube by using the NE-373-A-6/4.2 kit (New Era Enterprises). RDC-based intervector projection angle restraints related to D_a_ and R values of −6 and 0.35, respectively.

For unbound mRBM20_(513-621)_ the ^1^H distances were obtained using NOE crosspeaks from 3D ^15^N-HSQC-NOESY (120 ms mixing time), 3D aliphatic ^13^C-HSQC-NOESY (120 ms mixing time), 3D aromatic ^13^C-HSQC-NOESY (120 ms mixing time) and 2D ^1^H,^1^H-NOESY (120 ms mixing time) spectra. Protein dihedral angles were obtained by using TALOS-N (Shen and Bax, 2015) and SideR (Hansen et al., 2010a; Hansen et al., 2010b).

### Relaxation measurement

^15^N relaxation data were acquired at 298 K and a field strength of 700 MHz. {^1^H}-^15^N heteronuclear NOE spectra were recorded with and without 3 s of proton saturation. The resulting values represent the average and standard deviation of two independent measurements.

### Isothermal Titration Calorimetry

Measurements were performed on an iTC_200_ microcalorimeter (Malvern Panalytical) at 298 K with a stir rate of 600 rpm and recorded with high sensitivity. The samples were dialyzed overnight in 20 mM sodium phosphate (pH 6.5), 50 mM NaCl and 2.5 mM TCEP prior to ITC experiments, and the same dialysis buffer was used to dilute protein and RNA samples. The cell contained target concentrations of 40 or 80 μM RNA, with target syringe concentrations of 400 or 800 μM protein, respectively (details in **Supplementary Figure 2**). After an initial delay of 120 s, a first injection of 1 uL was followed by 12 injections of 3 uL, with a delay of 120 s between each injection. Data processing used NITPIC (Keller et al., 2012); (Scheuermann and Brautigam, 2015) to integrate the titration points, and SEDPHAT (Zhao et al., 2015a) to perform the curve fitting. The values in Table 1 present averages and standard deviations from at least two independent measurements. The graph in Fig. 1B was prepared by using GUSSI (Brautigam, 2015).

### Data deposition

The structure ensemble of mRBM20_(513-621)_ bound to AUCUUA RNA has been deposited at the Protein Data Bank (http://www.ebi.ac.uk/pdbe/) with accession ID 6SO9. The ensemble of structures for unbound mRBM20_(513-621)_ has also been deposited, with accession ID 6SOE. Chemical shift assignments for RNA-bound and unbound mRBM20_(513-621)_ have been deposited in the Biological Magnetic Resonance Data Bank (http://bmrb.wisc.edu/) under BMRB accession numbers 34428 and 34429, respectively.

## Acknowledgements

Plasmids encoding mouse RBM20 cDNA were provided by Pamela Lorenzi of the Department of Neurological, Biomedical and Movement Sciences, University of Verona. We thank Axelle Grélard, Estelle Morvan, and the structural biology platform at the Institut Europèen de Chimie et Biologie for access to the NMR spectrometers, equipment and technical assistance. Financial support from the IR-RMN-THC Fr3050 CNRS for conducting the research is gratefully acknowledged. SKU is supported by a DST INSPIRE Faculty Award (IFA-13 CH-102) from the Government of India Department of Science and Technology. SKU thanks Dr. Souvik Maiti for support and assistance with the research. CDM acknowledges financial support from INSERM, CNRS and University of Bordeaux.

## REFERENCES

Ashkenazy, H., Abadi, S., Martz, E., Chay, O., Mayrose, I., Pupko, T., and Ben-Tal, N. (2016). ConSurf 2016: an improved methodology to estimate and visualize evolutionary conservation in macromolecules. Nucleic Acids Res. 44, W344–W350.

Azad, A., Poloni, G., Sontayananon, N., Jiang, H., and Gehmlich, K. (2019). The giant titin: how to evaluate its role in cardiomyopathies. J. Muscle Res. Cell Motil. 40, 159–167.

Beraldi, R., Li, X., Martinez Fernandez, A., Reyes, S., Secreto, F., Terzic, A., Olson, T.M., and Nelson, T.J. (2014). Rbm20-deficient cardiogenesis reveals early disruption of RNA processing and sarcomere remodeling establishing a developmental etiology for dilated cardiomyopathy. Hum. Mol. Genet. 23, 3779–3791.

Berezin, C., Glaser, F., Rosenberg, J., Paz, I., Pupko, T., Fariselli, P., Casadio, R., and Ben-Tal, N. (2004). ConSeq: the identification of functionally and structurally important residues in protein sequences. Bioinformatics 20, 1322–1324.

Bertero, A., Fields, P.A., Ramani, V., Bonora, G., Yardimci, G.G., Reinecke, H., Pabon, L., Noble, W.S., Shendure, J., and Murry, C.E. (2019). Dynamics of genome reorganization during human cardiogenesis reveal an RBM20-dependent splicing factory. Nat. Commun. 10, 1538.

Brauch, K.M., Karst, M.L., Herron, K.J., de Andrade, M., Pellikka, P.A., Rodeheffer, R.J., Michels, V.V., and Olson, T.M. (2009). Mutations in ribonucleic acid binding protein gene cause familial dilated cardiomyopathy. J. Am. Coll. Cardiol. 54, 930–941.

Brautigam, C.A. (2015). Calculations and Publication-Quality Illustrations for Analytical Ultracentrifugation Data. Methods Enzymol. 562, 109–133.

Brunger, A.T. (2007). Version 1.2 of the Crystallography and NMR system. Nat. Protoc. 2, 2728–2733.

Buck, D., Smith, J.E., 3rd, Chung, C.S., Ono, Y., Sorimachi, H., Labeit, S., and Granzier, H.L. (2014). Removal of immunoglobulin-like domains from titin’s spring segment alters titin splicing in mouse skeletal muscle and causes myopathy. J. Gen. Physiol. 143, 215–230.

Bull, M., Methawasin, M., Strom, J., Nair, P., Hutchinson, K., and Granzier, H. (2016). Alternative Splicing of Titin Restores Diastolic Function in an HFpEF-Like Genetic Murine Model (TtnΔIAjxn). Circ. Res. 119, 764–772.

Cai, H., Zhu, C., Chen, Z., Maimaiti, R., Sun, M., McCormick, R.J., Lan, X., Chen, H., and Guo, W. (2019). Angiotensin II Influences Pre-mRNA Splicing Regulation by Enhancing RBM20 Transcription Through Activation of the MAPK/ELK1 Signaling Pathway. Int. J. Mol. Sci. 20, 5059.

Chami, N., Tadros, R., Lemarbre, F., Lo, K.S., Beaudoin, M., Robb, L., Labuda, D., Tardif, J.-C., Racine, N., Talajic, M., et al. (2014). Nonsense mutations in BAG3 are associated with early-onset dilated cardiomyopathy in French Canadians. Can. J. Cardiol. 30, 1655–1661.

Cornilescu, G., Marquardt, J.L., Ottiger, M., Bax, A. (1998). Validation of protein structure from anisotropic carbonyl chemical shifts in a dilute liquid crystalline phase. J. Am. Chem. Sci. 120, 6836–6837.

Dauksaite, V., and Gotthardt, M. (2018). Molecular basis of titin exon exclusion by RBM20 and the novel titin splice regulator PTB4. Nucleic Acids Res. 46, 5227–5238.

Delaglio, F., Grzesiek, S., Vuister, G., Zhu, G., Pfeifer, J., and Bax, A. (1995). NMRPipe: A multidimensional spectral processing system based on UNIX pipes. Journal of Biomolecular NMR 6.

Dorn, G., Leitner, A., Boudet, J., Campagne, S., von Schroetter, C., Moursy, A., Aebersold, R., and Allain, F.H.-T. (2017). Structural modeling of protein-RNA complexes using crosslinking of segmentally isotope-labeled RNA and MS/MS. Nat. Methods 14, 487–490.

Filippello, A., Lorenzi, P., Bergamo, E., and Romanelli, M.G. (2013). Identification of nuclear retention domains in the RBM20 protein. FEBS Lett. 587, 2989–2995.

Gigli, M., Begay, R.L., Morea, G., Graw, S.L., Sinagra, G., Taylor, M.R.G., Granzier, H., and Mestroni, L. (2016). A Review of the Giant Protein Titin in Clinical Molecular Diagnostics of Cardiomyopathies. Front. Cardiovasc. Med. 3, 21.

Granzier, H., and Labeit, S. (2002). Cardiac titin: an adjustable multi-functional spring. J. Physiol. 541, 335–342.

Greaser, M.L., Krzesinski, P.R., Warren, C.M., Kirkpatrick, B., Campbell, K.S., and Moss, R.L. (2005). Developmental changes in rat cardiac titin/connectin: transitions in normal animals and in mutants with a delayed pattern of isoform transition. J. Muscle Res. Cell Motil. 26, 325–332.

Guo, W., Bharmal, S.J., Esbona, K., and Greaser, M.L. (2010). Titin diversity--alternative splicing gone wild. J. Biomed. Biotechnol. 2010, 753675.

Guo, W., Schafer, S., Greaser, M.L., Radke, M.H., Liss, M., Govindarajan, T., Maatz, H., Schulz, H., Li, S., Parrish, A.M., et al. (2012). RBM20, a gene for hereditary cardiomyopathy, regulates titin splicing. Nat. Med. 18, 766–773.

Guo, W., Pleitner, J.M., Saupe, K.W., and Greaser, M.L. (2013). Pathophysiological defects and transcriptional profiling in the RBM20-/- rat model. PLoS One 8, e84281.

Guo, W., Zhu, C., Yin, Z., Wang, Q., Sun, M., Cao, H., and Greaser, M.L. (2018). Splicing Factor RBM20 Regulates Transcriptional Network of Titin Associated and Calcium Handling Genes in The Heart. Int. J. Biol. Sci. 14, 369–380.

Hansen, D.F., Neudecker, P., and Kay, L.E. (2010a). Determination of isoleucine side-chain conformations in ground and excited states of proteins from chemical shifts. J. Am. Chem. Soc. 132, 7589–7591.

Hansen, D.F., Neudecker, P., Vallurupalli, P., Mulder, F.A.A., and Kay, L.E. (2010b). Determination of Leu side-chain conformations in excited protein states by NMR relaxation dispersion. J. Am. Chem. Soc. 132, 42–43.

Hey, T.M., Rasmussen, T.B., Madsen, T., Aagaard, M.M., Harbo, M., Mølgaard, H., Møller, J.E., Eiskjær, H., and Mogensen, J. (2019). Pathogenic RBM20-Variants Are Associated With a Severe Disease Expression in Male Patients With Dilated Cardiomyopathy. Circ. Heart Fail. 12, e005700

Hinze, F., Dieterich, C., Radke, M.H., Granzier, H., and Gotthardt, M. (2016). Reducing RBM20 activity improves diastolic dysfunction and cardiac atrophy. J. Mol. Med. 94, 1349–1358.

van den Hoogenhof, M.M.G., Beqqali, A., Amin, A.S., van der Made, I., Aufiero, S., Khan, M.A.F., Schumacher, C.A., Jansweijer, J.A., van Spaendonck-Zwarts, K.Y., Remme, C.A., et al. (2018). RBM20 Mutations Induce an Arrhythmogenic Dilated Cardiomyopathy Related to Disturbed Calcium Handling. Circulation 138, 1330–1342.

Ito, J., Iijima, M., Yoshimoto, N., Niimi, T., Kuroda, S. ’ichi, and Maturana, A.D. (2016). RBM20 and RBM24 cooperatively promote the expression of short enh splice variants. FEBS Lett. 590, 2262–2274.

Kayvanpour, E., Sedaghat-Hamedani, F., Amr, A., Lai, A., Haas, J., Holzer, D.B., Frese, K.S., Keller, A., Jensen, K., Katus, H.A., et al. (2017). Genotype-phenotype associations in dilated cardiomyopathy: meta-analysis on more than 8000 individuals. Clin. Res. Cardiol. 106, 127–139.

Keller, S., Vargas, C., Zhao, H., Piszczek, G., Brautigam, C.A., and Schuck, P. (2012). High-precision isothermal titration calorimetry with automated peak-shape analysis. Anal. Chem. 84, 5066–5073.

Khan, M.A.F., Reckman, Y.J., Aufiero, S., van den Hoogenhof, M.M.G., van der Made, I., Beqqali, A., Koolbergen, D.R., Rasmussen, T.B., van der Velden, J., Creemers, E.E., et al. (2016). RBM20 Regulates Circular RNA Production From the Titin Gene. Circ. Res. 119, 996–1003.

Kimura, A. (2016). Molecular genetics and pathogenesis of cardiomyopathy. J. Hum. Genet. 61, 41–50.

Laskowski, R.A., Rullmannn, J.A., MacArthur, M.W., Kaptein, R., and Thornton, J.M. (1996). AQUA and PROCHECK-NMR: programs for checking the quality of protein structures solved by NMR. J. Biomol. NMR 8, 477–486.

Li, D., Morales, A., Gonzalez-Quintana, J., Norton, N., Siegfried, J.D., Hofmeyer, M., and Hershberger, R.E. (2010). Identification of novel mutations in RBM20 in patients with dilated cardiomyopathy. Clin. Transl. Sci. 3, 90–97.

Li, S., Guo, W., Dewey, C.N., and Greaser, M.L. (2013). Rbm20 regulates titin alternative splicing as a splicing repressor. Nucleic Acids Research 41, 2659–2672.

Liss, M., Radke, M.H., Eckhard, J., Neuenschwander, M., Dauksaite, V., von Kries, J.-P., and Gotthardt, M. (2018). Drug discovery with an RBM20 dependent titin splice reporter identifies cardenolides as lead structures to improve cardiac filling. PLoS One 13, e0198492.

Liu, Y., and Prestegard, J.H. (2010). A device for the measurement of residual chemical shift anisotropy and residual dipolar coupling in soluble and membrane-associated proteins. J. Biomol. NMR 47, 249–258.

Long, P.A., Evans, J.M., and Olson, T.M. (2017). Diagnostic Yield of Whole Exome Sequencing in Pediatric Dilated Cardiomyopathy. J. Cardiovasc. Dev. Dis. 4, E11

Lorenzi, P., Sangalli, A., Fochi, S., Dal Molin, A., Malerba, G., Zipeto, D., and Romanelli, M.G. (2019). RNA-binding proteins RBM20 and PTBP1 regulate the alternative splicing of FHOD3. Int. J. Biochem. Cell Biol. 106, 74–83.

Maatz, H., Jens, M., Liss, M., Schafer, S., Heinig, M., Kirchner, M., Adami, E., Rintisch, C., Dauksaite, V., Radke, M.H., et al. (2014). RNA-binding protein RBM20 represses splicing to orchestrate cardiac pre-mRNA processing. J. Clin. Invest. 124, 3419–3430.

Maiti, R., Van Domselaar, G.H., Zhang, H., and Wishart, D.S. (2004). SuperPose: a simple server for sophisticated structural superposition. Nucleic Acids Res. 32, W590–W594.

Maruyama, K., Natori, R., and Nonomura, Y. (1976). New elastic protein from muscle. Nature 262, 58–60.

Methawasin, M., Hutchinson, K.R., Lee, E.-J., Smith, J.E., 3rd, Saripalli, C., Hidalgo, C.G., Ottenheijm, C.A.C., and Granzier, H. (2014). Experimentally increasing titin compliance in a novel mouse model attenuates the Frank-Starling mechanism but has a beneficial effect on diastole. Circulation 129, 1924–1936.

Methawasin, M., Strom, J.G., Slater, R.E., Fernandez, V., Saripalli, C., and Granzier, H. (2016). Experimentally Increasing the Compliance of Titin Through RNA Binding Motif-20 (RBM20) Inhibition Improves Diastolic Function In a Mouse Model of Heart Failure With Preserved Ejection Fraction. Circulation 134, 1085–1099.

Millat, G., Bouvagnet, P., Chevalier, P., Sebbag, L., Dulac, A., Dauphin, C., Jouk, P.-S., Delrue, M.-A., Thambo, J.-B., Le Metayer, P., et al. (2011). Clinical and mutational spectrum in a cohort of 105 unrelated patients with dilated cardiomyopathy. Eur. J. Med. Genet. 54, e570–e575.

Murayama, R., Kimura-Asami, M., Togo-Ohno, M., Yamasaki-Kato, Y., Naruse, T.K., Yamamoto, T., Hayashi, T., Ai, T., Spoonamore, K.G., Kovacs, R.J., et al. (2018). Phosphorylation of the RSRSP stretch is critical for splicing regulation by RNA-Binding Motif Protein 20 (RBM20) through nuclear localization. Sci. Rep. 8, 8970.

Nielsen, J.B., Thorolfsdottir, R.B., Fritsche, L.G., Zhou, W., Skov, M.W., Graham, S.E., Herron, T.J., McCarthy, S., Schmidt, E.M., Sveinbjornsson, G., et al. (2018). Biobank-driven genomic discovery yields new insight into atrial fibrillation biology. Nat. Genet. 50, 1234–1239.

Oberstrass, F.C., Auweter, S.D., Erat, M., Hargous, Y., Henning, A., Wenter, P., Reymond, L., Amir-Ahmady, B., Pitsch, S., Black, D.L., et al. (2005). Structure of PTB bound to RNA: specific binding and implications for splicing regulation. Science 309, 2054–2057.

Parikh, V.N., Caleshu, C., Reuter, C., Lazzeroni, L.C., Ingles, J., Garcia, J., McCaleb, K., Adesiyun, T., Sedaghat-Hamedani, F., Kumar, S., et al. (2019). Regional Variation in RBM20 Causes a Highly Penetrant Arrhythmogenic Cardiomyopathy. Circ. Heart Fail. 12, e005371.

van der Pijl, R., Strom, J., Conijn, S., Lindqvist, J., Labeit, S., Granzier, H., and Ottenheijm, C. (2018). Titin-based mechanosensing modulates muscle hypertrophy. J. Cachexia Sarcopenia Muscle 9, 947–961.

Pulcastro, H.C., Awinda, P.O., Methawasin, M., Granzier, H., Dong, W., and Tanner, B.C.W. (2016). Increased Titin Compliance Reduced Length-Dependent Contraction and Slowed Cross-Bridge Kinetics in Skinned Myocardial Strips from Rbm (20ΔRRM) Mice. Front. Physiol. 7, 322.

Rampersaud, E., Siegfried, J.D., Norton, N., Li, D., Martin, E., and Hershberger, R.E. (2011). Rare variant mutations identified in pediatric patients with dilated cardiomyopathy. Prog. Pediatr. Cardiol. 31, 39–47.

Refaat, M.M., Lubitz, S.A., Makino, S., Islam, Z., Frangiskakis, J.M., Mehdi, H., Gutmann, R., Zhang, M.L., Bloom, H.L., MacRae, C.A., et al. (2012). Genetic variation in the alternative splicing regulator RBM20 is associated with dilated cardiomyopathy. Heart Rhythm 9, 390–396.

Rieping, W., Habeck, M., Bardiaux, B., Bernard, A., Malliavin, T.E., and Nilges, M. (2007). ARIA2: automated NOE assignment and data integration in NMR structure calculation. Bioinformatics 23, 381–382.

Rindler, T.N., Hinton, R.B., Salomonis, N., and Ware, S.M. (2017). Molecular Characterization of Pediatric Restrictive Cardiomyopathy from Integrative Genomics. Sci. Rep. 7, 39276.

Robyns, T., Willems, R., Van Cleemput, J., Jhangiani, S., Muzny, D., Gibbs, R., Lupski, J.R., Breckpot, J., Devriendt, K., and Corveleyn, A. (2019). Whole exome sequencing in a large pedigree with DCM identifies a novel mutation in RBM20. Acta Cardiologica 1–6.

Scheuermann, T.H., and Brautigam, C.A. (2015). High-precision, automated integration of multiple isothermal titration calorimetric thermograms: new features of NITPIC. Methods 76, 87–98.

Sedaghat-Hamedani, F., Haas, J., Zhu, F., Geier, C., Kayvanpour, E., Liss, M., Lai, A., Frese, K., Pribe-Wolferts, R., Amr, A., et al. (2017). Clinical genetics and outcome of left ventricular non-compaction cardiomyopathy. Eur. Heart J. 38, 3449–3460.

Senn, H., Werner, B., Messerle, B.A., Weber, C., Traber, R., and Wüthrich, K. (1989). Stereospecific assignment of the methyl 1 H NMR lines of valine and leucine in polypeptides by nonrandom 13 C labelling. FEBS Lett. 249, 113–118.

Shen, Y., and Bax, A. (2015). Protein structural information derived from NMR chemical shift with the neural network program TALOS-N. Methods Mol. Biol. 1260, 17–32.

Tamiola, K., Acar, B., and Mulder, F.A.A. (2010). Sequence-specific random coil chemical shifts of intrinsically disordered proteins. J. Am. Chem. Soc. 132, 18000–18003.

Waldmüller, S., Schroeder, C., Sturm, M., Scheffold, T., Imbrich, K., Junker, S., Frische, C., Hofbeck, M., Bauer, P., Bonin, M., et al. (2015). Targeted 46-gene and clinical exome sequencing for mutations causing cardiomyopathies. Mol. Cell. Probes 29, 308–314.

Wang, K., McClure, J., and Tu, A. (1979). Titin: major myofibrillar components of striated muscle. Proc. Natl. Acad. Sci. U S A 76, 3698–3702.

Watanabe, T., Kimura, A., and Kuroyanagi, H. (2018). Alternative Splicing Regulator RBM20 and Cardiomyopathy. Front. Mol. Biosci. 5, 105.

Wells, Q.S., Becker, J.R., Su, Y.R., Mosley, J.D., Weeke, P., D’Aoust, L., Ausborn, N.L., Ramirez, A.H., Pfotenhauer, J.P., Naftilan, A.J., et al. (2013). Whole exome sequencing identifies a causal RBM20 mutation in a large pedigree with familial dilated cardiomyopathy. Circ. Cardiovasc. Genet. 6, 317–326.

Wyles, S.P., Li, X., Hrstka, S.C., Reyes, S., Oommen, S., Beraldi, R., Edwards, J., Terzic, A., Olson, T.M., and Nelson, T.J. (2016). Modeling structural and functional deficiencies of RBM20 familial dilated cardiomyopathy using human induced pluripotent stem cells. Hum. Mol. Genet. 25, 254–265.

Yachdav, G., Kloppmann, E., Kajan, L., Hecht, M., Goldberg, T., Hamp, T., Hönigschmid, P., Schafferhans, A., Roos, M., Bernhofer, M., et al. (2014). PredictProtein--an open resource for online prediction of protein structural and functional features. Nucleic Acids Res. 42, W337–W343.

Yamamoto, T., Miura, A., Itoh, K., Takeshima, Y., and Nishio, H. (2019). RNA sequencing reveals abnormal LDB3 splicing in sudden cardiac death. Forensic Sci. Int. 302, 109906.

Zhao, H., Piszczek, G., and Schuck, P. (2015a). SEDPHAT – A platform for global ITC analysis and global multi-method analysis of molecular interactions. Methods 76, 137–148.

Zhao, Y., Feng, Y., Zhang, Y.-M., Ding, X.-X., Song, Y.-Z., Zhang, A.-M., Liu, L., Zhang, H., Ding, J.-H., and Xia, X.-S. (2015b). Targeted next-generation sequencing of candidate genes reveals novel mutations in patients with dilated cardiomyopathy. Int. J. Mol. Med. 36, 1479–1486.

Zhu, C., Yin, Z., Ren, J., McCormick, R.J., Ford, S.P., and Guo, W. (2015). RBM20 is an essential factor for thyroid hormone-regulated titin isoform transition. J. Mol. Cell Biol. 7, 88–90.

Zhu, C., Yin, Z., Tan, B., and Guo, W. (2017). Insulin regulates titin pre-mRNA splicing through the PI3K-Akt-mTOR kinase axis in a RBM20-dependent manner. Biochim. Biophys. Acta Mol. Basis Dis. 1863, 2363–2371.

