## Supplementary Material for "Structural basis of UCUU RNA motif recognition by splicing factor RBM20"

Santosh Kumar Upadhyay, Cameron D. Mackereth

SUPPLEMENTARY MATERIAL

**Main Resources**

**Supplementary Table 1.** List of PCR primers.

**Supplementary Figure 1.** Annotated 2D $^{1}H,^{15}N$-HSQC spectra corresponding to **Fig. 1C**.

**Supplementary Figure 2.** Representative ITC raw data corresponding to **Fig. 1B** and **Table 1**.

**Supplementary Figure 3.** Complete spectra corresponding to the selected regions in **Fig. 3C-F**.

### Main Resources

| REAGENT or RESOURCE | SOURCE | IDENTIFIER |
|---|---|---|
| Bacterial and Virus Strains | | |
| 5-alpha Competent E. coli | New England Biolabs | Cat# C2987H |
| T7 Express lysY Competent E. coli | New England Biolabs | Cat# C3010I |
| Chemicals, Peptides, and Recombinant Proteins | | |
| Nuvia™ IMAC Resin | BioRad | Cat# 7800800 |
| TEV protease | In house produced | N/A |
| Deposited Data | | |
| RBM20-AUCUUA chemical shifts | BioMagResBank (BMRB) | 34428 |
| RBM20-AUCUUA structural ensemble | Protein Data Bank (PDB) | 6SO9 |
| unbound RBM20 chemical shifts | BioMagResBank (BMRB) | 34429 |
| unbound RBM20 structural ensemble | Protein Data Bank (PDB) | 6SOE |
| Oligonucleotides | | |
| PCR oligos | Sigma | Supplementary Table 1 |
| RNA | In house produced | Table 1 |
| Recombinant DNA | | |
| pET-His1a | EMBL | N/A |
| mouse RBM20 cDNA | Pamela Lorenzi, University of Verona | N/A |
| pET-His1a:mRBM20 (513-621)H523A | This paper | N/A |
| pET-His1a:mRBM20 (513-621)N526A | This paper | N/A |
| pET-His1a:mRBM20 (513-621)V537I | This paper | N/A |
| pET-His1a:mRBM20 (513-621)Q558A | This paper | N/A |
| pET-His1a:mRBM20 (513-621)F560A | This paper | N/A |
| pET-His1a:mRBM20 (513-621)Q577A | This paper | N/A |
| pET-His1a:mRBM20 (513-621)R591M | This paper | N/A |
| pET-His1a:mRBM20 (513-621)R595M | This paper | N/A |
| pET-His1a:mRBM20 (513-621)Y596A | This paper | N/A |
| pET-His1a:mRBM20 (513-609)Δα3 | This paper | N/A |
| pET-His1a:mRBM20 (513-649)+RS | This paper | N/A |
| Software and Algorithms | | |
| Topspin | Bruker BioSpin | Version 4.0 |
| NMRPipe | Delaglio et al, 1995 | Version 8.6 |
| Sparky | T.D. Goddard and D.G. Kneller, Univ. of California | Version 3 |
| CNS | | Version 1.2 |
| ARIA | | Version 2.3 |

**Supplementary Table 1.** List of PCR primers.

| Function | Direction | Primer sequence (5'→3') |
|---|---|---|
| Clone mRBM20 (513-621) from cDNA, add stop codon, *Nco*I/*Acc65*I sites | Forward | CAGTAGCCATGGCACAGAGGAAAGGCGCTG |
| | Reverse | GTGGTACCTTACCTCTCCCTCTGGGAATGG |
| Remove C-terminal helix (-609), add stop codon, *Acc65*I site | Reverse | GTGGTACCTTAAGCCACATTTTTCCCAGGTTTC |
| Add RS region (-649), add stop codon, *Acc65I* site | Reverse | GTGGTACCTTAGGATCTTGGGGAGAGTGATC |
| H523A mutation | Forward | CGGGTAGTGGCGATCTGCAATCTCCCG |
| | Reverse | GATTGCAGATCGCCACTACCCGTCCAGC |
| N526A mutation | Forward | CACATCTGCGCGCTCCCGGAGGGCAGC |
| | Reverse | GCCCTCCGGGAGCGCGCAGATGTGCACTAC |
| Q558A mutation | Forward | GTCAACTAATGCGGCTTTCTTGGAG |
| | Reverse | CAAGAAAGCCGCATTAGTTGACTTCATGAG |
| V537I mutation | Forward | GAGAATGACATTATTAACCTGGGGCTGCCC |
| | Reverse | CAGGTTAATAATGTCATTCTCCGTGCAGC |
| F560A mutation | Forward | CTAATCAGGCTGCGTTGGAGATGGCTTAC |
| | Reverse | GCCATCTCCAACGCAGCCTGATTAGTTGACTTC |
| Q577A mutation | Forward | CAGTACTACGCGGAAAAGCCTGCGATTATC |
| | Reverse | GCAGGCTTTTCCGCGTAGTACTGGACCATAGC |
| K779M mutation | Forward | CTACCAAGAAATGCCTGCGATTATCAATG |
| | Reverse | GATAATCGCAGGCATTTCTTGGTAGTACTG |
| R591M mutation | Forward | GTTACTCATTATGATGTCCACCAGATACAAG |
| | Reverse | CTGGTGGACATCATAATGAGTAACTTCTCGCC |
| R595M mutation | Forward | CATGTCCACCATGTACAAGGAATTGCAGC |
| | Reverse | CCTTGTACATGGTGGACATGCGAATGAG |
| Y596A mutation | Forward | GTCCACCAGAGCGAAGGAATTGCAGCTG |
| | Reverse | GCAATTCCTTCGCTCTGGTGGACATGCG |

A)

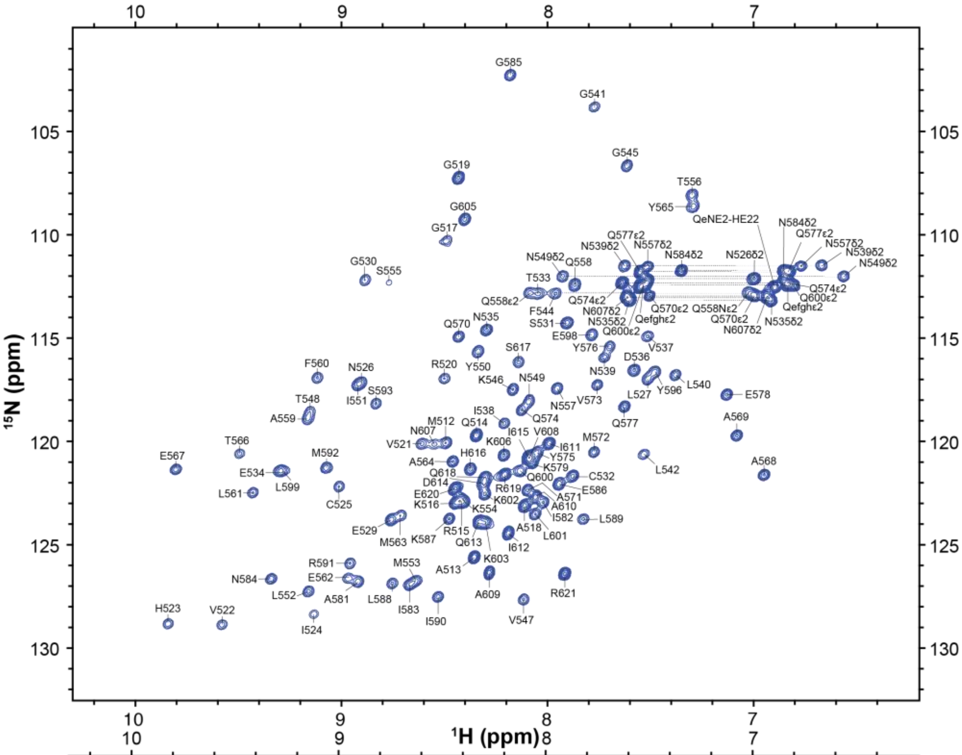


B)

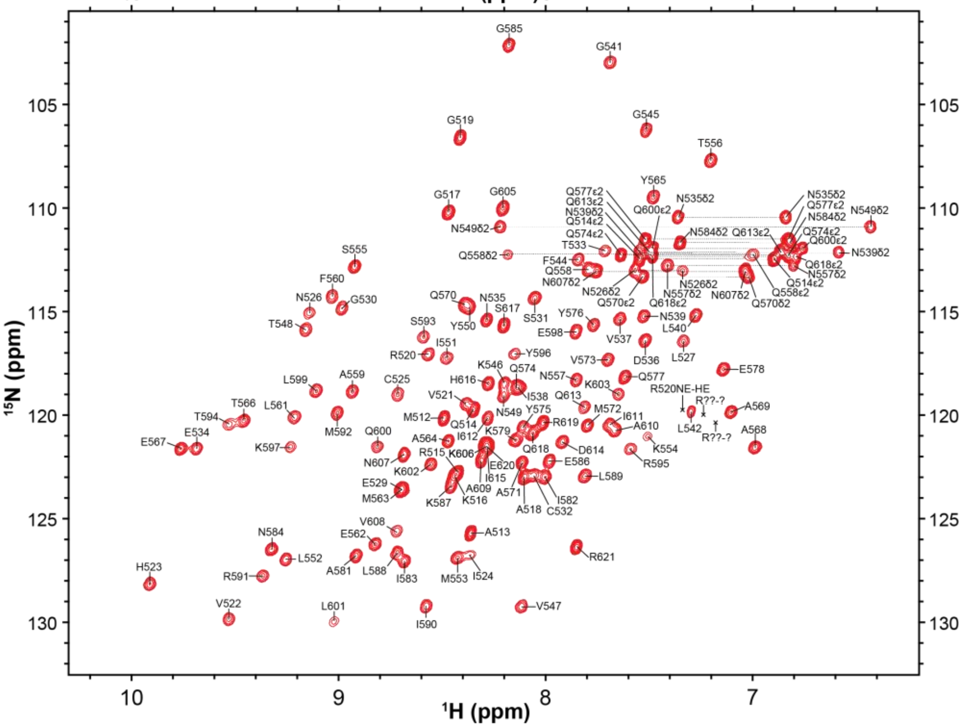


**Supplementary Figure 1.** Annotated 2D $^{1}H,^{15}N$-HSQC spectra corresponding to **Fig. 1C**. The backbone and side chain amide cross peaks are labeled with the residue name (single amino acid letter code) and number. The spectra were recorded at 298 K in 20 mM sodium phosphate, 50 mM NaCl (pH 6.5), 2 mM DTT and 10 % (v/v) $D_2O$. (**A**) NMR sample containing 400 µM [$^{13}C,^{15}N$]mRBM20$_{(513\text{-}621)}$ and spectrum collected at a field strength of 800 MHz. (**B**) NMR sample containing 400 µM [$^{13}C,^{15}N$]mRBM20$_{(513\text{-}621)}$ and 480 µM natural abundance AUCUUA RNA, with spectrum collected at a field strength of 700 MHz.

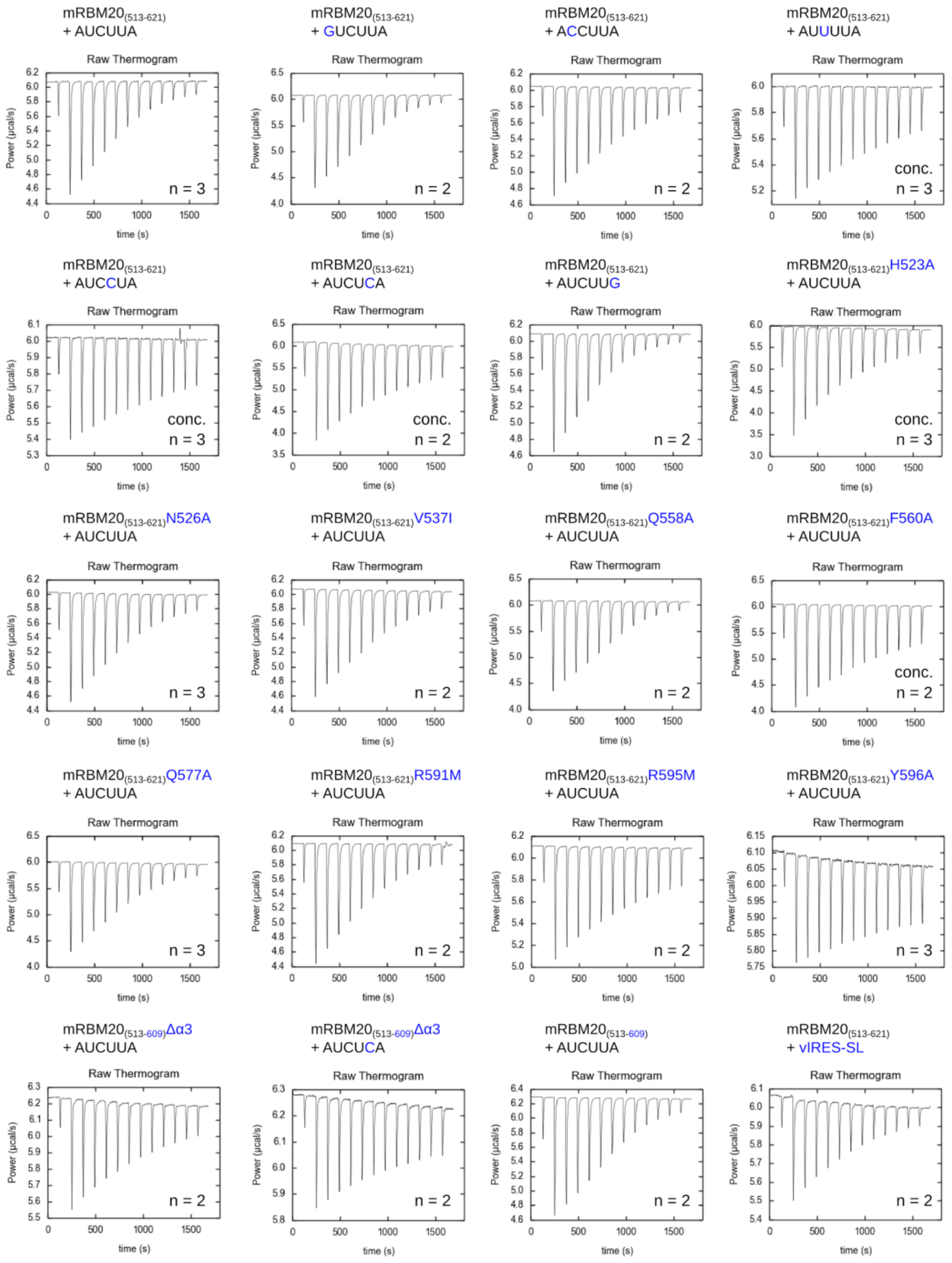


**Supplementary Figure 2.** Representative ITC raw data corresponding to **Fig. 1B** and **Table 1**. Above each thermogram the protein construct in the syringe is listed first, followed by the RNA ligand present in the cell. The number of experiments used to calculate the average and standard deviation in **Table 1** is indicated at the bottom right of each thermogram. In most cases, the target protein concentration was 400 µM, with 40 µM RNA ligand. Those which required higher concentrations of 800 µM protein and 80 µM RNA are indicated with ‘conc.’.

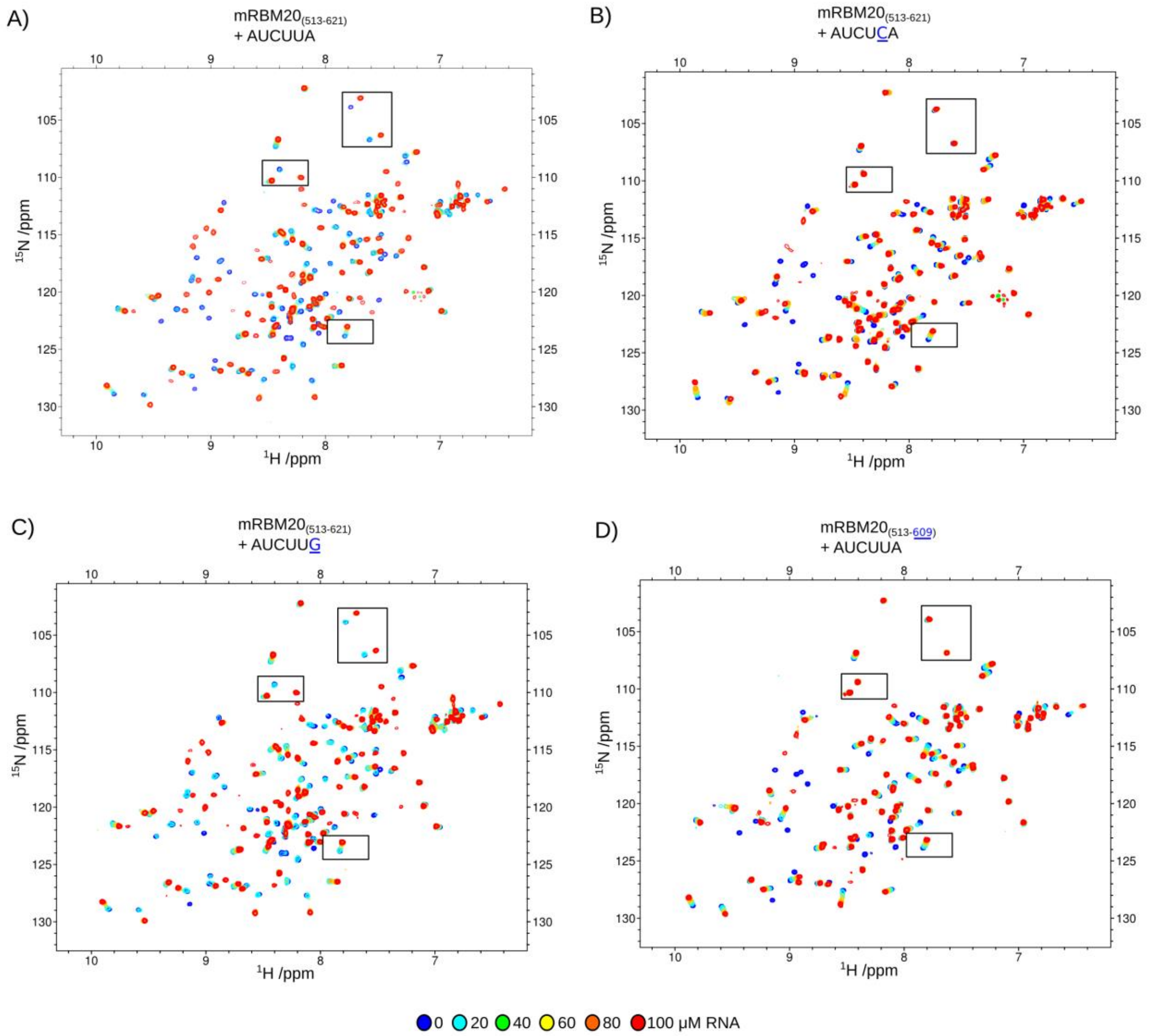


**Supplementary Figure 3.** Complete spectra corresponding to the selected regions in **Fig. 3C-F**.
